# How interactions between temperature and resources scale from populations to communities

**DOI:** 10.1101/2024.09.19.613936

**Authors:** Colin T. Kremer, Mridul K. Thomas, Christopher A. Klausmeier, Elena Litchman

## Abstract

Temperature and resources are fundamental factors that determine the ability of organisms to function and survive, while influencing their development, growth, and reproduction. Major bodies of ecological theory have emerged, largely independently, to address temperature and resource effects. It remains a major challenge to unite these ideas and determine the interactive effects of temperature and resources on ecological patterns and processes, and their consequences across ecological scales. Here, we propose a simple, physiologically motivated model capturing the interactive effects of temperature and resources (including inorganic nutrients and light) on the growth of microbial ectotherms over multiple ecological scales. From this model we derive a set of key predictions. At the population level, we predict (i) interactive effects of resource limitation on thermal traits, (ii) consistent differences in the temperature sensitivity of auto- and heterotrophs, and (iii) the existence of specific tradeoffs between traits that determine the shape of thermal performance curves. At the community level, we derive predictions for (iv) how limitation by nutrients and light can change the relationship between temperature and productivity. All four predictions are upheld, based on our analyses of a large compilation of laboratory data on microbial growth, as well as field experiments with marine phytoplankton communities. Collectively, our modeling framework provides a new way of thinking about the interplay between two fundamental aspects of life — temperature and resources — and how they constrain and structure ecological properties across scales. Providing links between population and community responses to simultaneous changes in abiotic factors is essential to anticipating the multifaceted effects of global change.

## Introduction

Understanding and predicting the ecological consequences of ongoing, multi-dimensional environmental change requires us to grapple with two major problems. First, while the effects of individual environmental drivers (temperature, pH, nutrients, food, etc.) are often reasonably well studied, interactions between drivers are often complex, difficult to measure, and poorly captured by existing theory and models. Second, it is exceedingly difficult to develop models that accurately capture reality at multiple scales (physiological, population, community), even for relatively simple processes such as growth and production. We address both of these problems, focusing on the effects of temperature-resource interactions on ectotherms, especially microorganisms. We propose a new modeling framework that applies to both population and community levels, and test model predictions across empirical data at multiple scales. Our results provide a foundation for exploring the biological effects of complex environmental changes using physiologically grounded models.

Resources and temperature are among the most fundamental factors regulating all life. Extensive studies, both empirical and theoretical, have investigated the effects of resources, especially inorganic nutrients, on individual growth, population dynamics, competition between species (Tilman 1980, Tilman et al. 1981, Grover 1997, Kooijman 2000, Nisbet et al. 2000), and ecosystem processes(Dillon and Rigler 1974, Sterner and Elser 2003). Meanwhile, temperature’s effects emerge chiefly from its strong influence on the rate of biochemical reactions.

Temperature is especially important for ectothermic organisms, which cannot regulate their internal body temperature (Angilletta 2009) and represent the majority of living things. At the level of individuals and populations, studies explore the effects of temperature on key traits, including development, growth, survival, reproduction, and fitness; such relationships are termed thermal performance curves (TPCs). At higher levels of organization, scaling relationships connecting metabolic rate to temperature are a major focus of the Metabolic Theory of Ecology (MTE) (Gillooly et al. 2001, Brown et al. 2004). Among other things, MTE predicts that the metabolic rate of heterotrophs increases more steeply with temperature than in photoautotrophs (Allen et al. 2005). This prediction is attributed to differences in activation energy of the rate-limiting biochemical processes that provide energy via respiration (carbohydrate metabolism, 0.65 eV) and photosynthesis (Rubisco, 0.32 eV), and is empirically supported (Brown et al. 2004, Allen et al. 2005, López-Urrutia et al. 2006, Kremer et al. 2017).

Studying interactions between abiotic factors — including temperature and resources — is difficult, especially with enough resolution and mechanistic detail to enable predictions of their ecological effects. Challenges arise because interactions can be complex and nonlinear, such that quantifying them may require large, logistically challenging experiments (Simmons et al. 2021, Pirotta et al. 2022; but see Thomas and Ranjan 2024). Regarding resources and temperature, at the level of individuals and populations, relatively recent results suggest a general pattern where resource limitation reduces the thermal tolerance and decreases the optimum temperature of organisms, including phytoplankton (Kremer et al. 2017, Bestion et al. 2018, Litchman and Thomas 2023), protists (Kimmance et al. 2006), zooplankton (Rueda Moreno and Sasaki 2023), insects (Huey and Kingsolver 2019, Huxley et al. 2021), and fish (Brett 1971). There are certainly many models accounting for the effects of multiple abiotic factors, including temperature and inorganic nutrients (Tilman et al. 1981, Lewington-Pearce et al. 2019). However, the way in which these effects are combined is often driven by mathematical convenience rather than empirically based relationships or detailed physiology (Thomas et al. 2017). Among other things, overlooking these important interactions changes predictions of species’ distributions and responses to climate change (Thomas et al. 2017). A growing number of studies exposing entire communities to combinations of temperature and resources also find important interactive effects (Ainsworth and Long 2021, Happe et al. 2024), although these outcomes are difficult to generalize and seldom predicted based on lower-level knowledge of the responses of individual community members. Without sufficiently mechanistic foundations, models of community and ecosystem processes rely on assumptions about the effects of interactions that may not reflect biological constraints and realities faced by individuals and populations (Pirotta et al. 2022).

A second major challenge in ecology is understanding how patterns and processes are connected across different scales of organization, from individuals to populations, communities, and ecosystems (Levin 1992, Chave 2013, Wickman et al. 2024). While a great many theories and models exist that are suited to making predictions at a particular scale, there are relatively few frameworks that successfully bridge multiple levels. Those that have emerged generally fall into two categories. Some capitalize on ever-expanding computational resources to create large, complex, and detailed models of low-level processes from which high-level patterns may emerge, such as individual based models (IBMs; Grimm et al. 2005, Railsback et al. 2020), or earth systems models resolving marine microbial diversity and biogeography (Follows et al. 2007) or terrestrial vegetation dynamics (Medvigy et al. 2010). Others derive simplifying approximations from underlying principles that capture major patterns spanning organizational scales, such as MTE’s exploration of temperature and size scaling relationships (Brown et al. 2004) or ecological stoichiometry’s focus on elemental ratios (Sterner and Elser 2003).

Accompanying their successes, both approaches also have limitations. Computational models are, and will continue to be, limited by computational power and depend on many poorly quantified parameters. Out of necessity, this can lead modelers to invoke untested assumptions and/or extensively tune models to match observations, which can lead to overfitting and therefore poor predictive power. Due to their complexity, these models may not actually advance our understanding even if they improve our prediction (which is itself uncertain). Conversely, scaling laws and approximations may often miss out on many important and biologically relevant details such as variation among taxonomic groups or around scaling relationships.

Notably, however, (Savage et al. 2004) elegantly derived connections between individual metabolism, resource supply, and population-level properties including growth rate and carrying capacity. Developing frameworks that span multiple scales is increasingly critical, as we struggle to predict the effects of anthropogenic environmental change, which are ultimately experienced by individuals, on regional and global processes.

Microbial systems, including phytoplankton (microbial photoautotrophs) are ideal for addressing these dual challenges of temperature-resource interactions and scale, due to their experimental tractability, extensive existing data, and ecological importance. The study of microorganisms and their ecophysiology is often comparatively simpler than for organisms with more complex behavior and life history. Growth depends primarily on temperature, energy (from light and/or organic matter), inorganic nutrients, and water. Individually, relationships between these abiotic factors and *population* growth are reasonably well-established for many organisms (e.g., for phytoplankton see Litchman and Klausmeier 2008). Studies of the effects of temperature on microbial growth go back more than 100 years (Dallinger 1887). These include extensive investigations of the thermal performance curves (TPCs) that relate population growth rate to environmental temperature, in both phytoplankton (Eppley 1972, Thomas et al. 2012, 2016, Boyd et al. 2013) and heterotrophic microbes (Corkrey et al. 2016). These TPCs are typically left-skewed unimodal curves with shapes that are well described by a variety of competing models (e.g., Grimaud et al. 2017, Padfield et al. 2021). Resource uptake and utilization has also been thoroughly investigated (e.g. Edwards et al. 2012), including an increasing number of studies investigating interactions between temperature and resources (reviewed in Litchman and Thomas 2023). Complementing these syntheses are a number of detailed physiological models (e.g. Geider et al. 1998, Flynn 2001). However, advances in linking these various physiological responses to population-level growth have not as readily produced general understanding of *community* growth rates. Yet understanding the rate at which both populations and communities grow is incredibly important. Individual species can shape ecosystems locally and globally, especially those that provide critical functions (e.g., nitrogen fixation by the marine phytoplankton *Trichodesmium* sp.) or have harmful effects (e.g., causing disease or producing toxins). The collective activity of entire microbial communities is also fundamentally important, driving global nutrient and energy cycles, including both fixing carbon via photosynthesis and liberating carbon through respiration.

Consequently, understanding the interactive effects of abiotic conditions on bulk community-level properties is essential in order to address questions about ecosystem- or global-scale processes. For example, how does primary production change across environmental gradients or in response to global change? Forecasts of atmospheric carbon, global temperatures and ocean biogeochemistry rely on accurately quantifying such relationships. Pragmatic solutions will prioritize identifying broad, general patterns without having to resolve in great detail the underlying complexities of populations or their interactions within communities. A classic example of such a relationship is the Eppley curve (Eppley 1972), an exponential function that empirically describes how the maximum possible phytoplankton growth rate across species increases with temperature (Bissinger et al. 2008, Kremer et al. 2017, Anderson et al. 2021). This ‘envelope function’ provides a community-level upper bound for all population-level TPCs and is a reasonable approximation for the functioning of diverse communities (Moisan et al. 2002); see also (Norberg et al. 2001, Thomas et al. 2012). The Eppley curve has enabled the creation of global ecosystem models incorporating flexible, simplified characterizations of ecophysiological processes and has been widely implemented to account for temperature sensitivity in phytoplankton (Moisan et al. 2002, Kremer et al. 2017, Anderson et al. 2024).

However, multiple questions about this important relationship remain. Regarding scaling, these include: How does the Eppley curve arise from underlying unimodal species-level TPCs? Also, what mechanisms control the steepness of this temperature-driven relationship (Bissinger et al. 2008, Kremer et al. 2017, Anderson et al. 2021)? Turning to temperature-resource interactions, how is the shape of this relationship modified in the presence of resource limitation? Answering these questions would improve our ability to effectively and efficiently predict the consequences of changing abiotic conditions.

Here we present a simple, physiologically motivated model of the effects of multiple abiotic factors including resources and temperature on population growth (Section 1). We then analyze a series of predictions arising from this model, evaluating them against experimental and observational data whenever possible (Section 2). This includes exploring the effects of multiple resource limitation on thermal responses (2.1), showing that the temperature-dependent growth of microorganisms with contrasting metabolic strategies agrees with predictions from Metabolic Theory (2.2), and identifying a trade-off constraining thermal performance (2.3). Finally, in Section 3 we show that, given certain assumptions, our population-level model can be extended to understand how growth is affected by temperature and resources at the community level.

Specifically, we show that the model both captures classic empirical relationships (the Eppley curve) and offers new insights into how these relationships are modified by resource availability, which are consistent with independent observational data. We conclude that our physiologically motivated model offers a valuable, intermediate-complexity approach to understanding the interactive effects of multiple important abiotic factors on the ecology of population and communities, with implications for ecosystem responses to climate change.

## 1. A population-level model of temperature- and resource-dependent growth

Population growth rate responds to multiple environmental drivers, many of which interact in complex ways. Modeling how populations and communities respond to realistic environmental variation requires us to specify mathematically how these drivers interact. Here, we first describe a model presented previously that captures how the growth rate of ectotherms depends on temperature and one resource (Thomas et al. 2017), then extend it to include multiple resources.

Population growth rate *μ* is the difference between per capita birth and death rates, which each depend on abiotic factors. Here, we assume birth and death rates increase exponentially with temperature due to thermodynamics and that resource limitation affects only the birth rate. The result is an equation for a typical left-skewed thermal performance curve (or TPC), describing how population growth rate varies with temperature (Logan et al. 1976, Thomas et al. 2017)

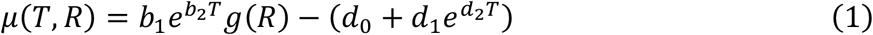

The first term of (1) represents birth rate, whose exponential increase with temperature *T* is controlled by the exponent *b*_2_, and also limited by resources *R* according to *g*(*R*), which ranges from zero to one. The second term of (1) accounts for temperature-independent (*d*_0_) and temperature-dependent 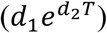 death. This definition of the model (1) clearly emphasizes our assumptions about the form and differences of temperature-dependence in birth and death. To a first approximation, these might also be interpreted as the difference between photosynthesis and respiration (Padfield et al. 2016, Schaum et al. 2017), although connecting these directly to net population growth rate has proved challenging, potentially requiring considering differences in carbon use efficiency (García-Carreras et al. 2018).

While the form of (1) is intuitive, we note that it can also be converted algebraically into an equivalent expression that is contains an explicit parameter for 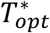 (the temperature at which growth rate is maximized when resources are not limiting) (O’Donnell et al. 2018). The result, (2), is useful in subsequent analyses:

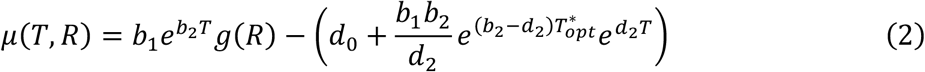

(See Appendix S2: Section S2 for a derivation, as well as a third version containing an explicit parameter for maximum growth rate). Here, we distinguish a population’s idealized optimum temperature, 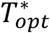, which is achieved only under optimal conditions (such as no resource limitation) from its realized *T*_*opt*_*(R)* under sub-optimal resource conditions, the maximum of *μ*(*T, R*) with respect to *T*. For example, (Thomas et al. 2017) show that 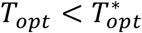 as resources decline.

To incorporate resource limitation of births, we define *g*(*R*) as a function that varies between zero and one as the availability of one or more resources *R* range from limiting to optimal. Where Thomas et al. (2017) focused on a single resource, we now allow birth to be limited by multiple resources (*R* = {*R*_1_, *R*_2_, … }). The interaction between resources can be modeled various ways, but as we lack sufficient data from a broad group of organisms to create a suitable mechanistic expression, we instead invoke Liebig’s law of the minimum (Liebig 1840) here, such that *g(R)* = min[*g*_1_(*R*_1_), *g*_2_(*R*_2_), …]. When limiting, each individual resource can affect birth rate according to its own functional form, *g*_i_(*R*_i_).

This general model can be specifically adapted to consider the growth of photoautotrophs, where birth is (co)-limited by both an inorganic nutrient *N* such as nitrogen or phosphorus and light *L*, yielding

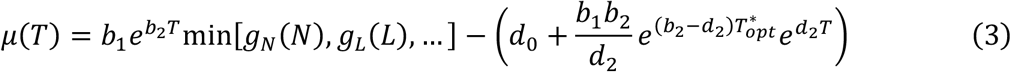

We further assume that nutrient limitation is governed by

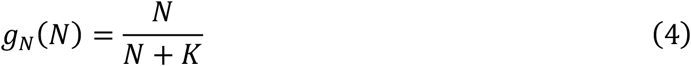

which is an increasing monotonic function of *N* with half-saturation constant *K* (**Fig. 1b**). Light limitation is more commonly described with a right-skewed unimodal function (as high light can cause photoinhibition), such as the Eilers-Peeters relationship (Eilers and Peeters 1988). We use a modified version of this relationship (Appendix S2: Section S3; **Fig. 1a**) to capture limitation due to light *L*, denoted

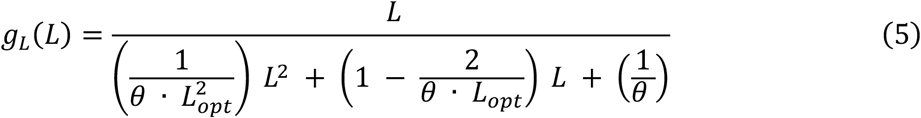

When light is optimal (*L* = *L*_*opt*_) this function achieves its maximum value of one. The shape parameter *θ* affects how quickly birth rates increase from low light levels (see Appendix S2: Section S3) and also how quickly they decrease above *L*_*opt*_.

**Figure 1.**
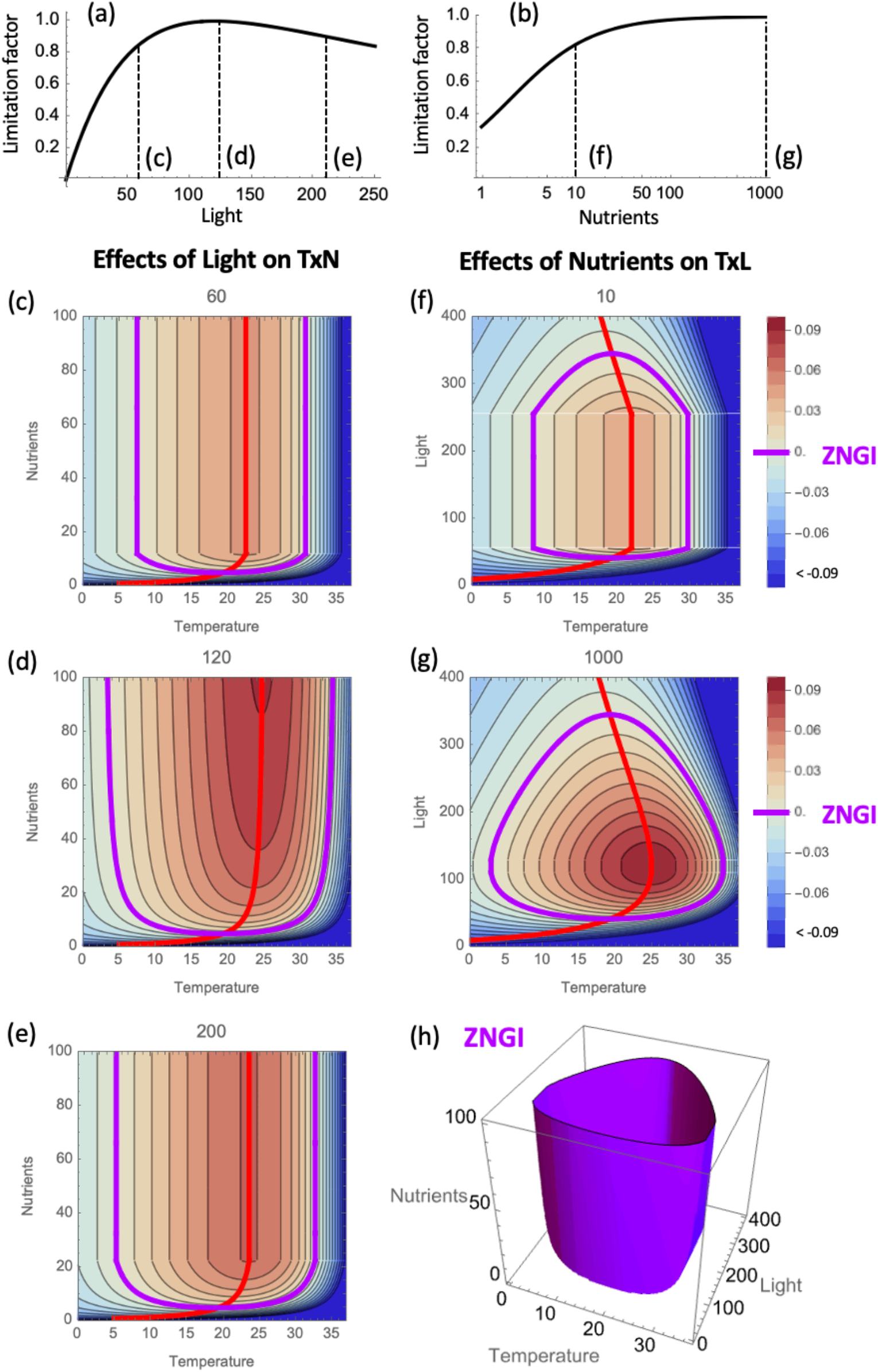
Both light (a) and nutrient availability (b) can impose limitations on growth rate. These effects can be factored into the temperature dependence of growth, as in (3), leading to complex interactions between temperature and nutrients (TxN) at different light levels (c-e) and between temperature and light (TxL) at different nutrient levels (f, g). In each panel, growth rate is indicated by shading and the combination of conditions leading to the zero net growth rate (ZNGI) is shown by the thick purple line. This forms a 3-dimensional surface (h) across conditions. Finally, the thick red lines in (c-g) illustrate how *T*_*opt*_ changes with nutrient and light availability. Model parameters are provided in Appendix S1: Table S1.

## 2. Analysis of population-level predictions

### 2.1 Effects of resource limitation

Several important predictions about population growth arise from our model (3): (i) *T*_*opt*_ declines with nutrient limitation (**Fig. 1d**), although this effect is most noticeable when light is not limiting (**Fig. 1 c,e**); (ii) *T*_*opt*_ declines as light levels deviate from optimal, leading to a unimodal relationship (**Fig. 1 f,g**), although this effect can be partly suppressed if nutrients are limiting (**Fig. 1f**); and (iii) the zero net growth isocline (ZNGI) also depends on all three abiotic factors, reproducing the U-shape characteristic of previous theory on temperature and a single resource (**Fig. 1 c-e**; Tilman et al. 1981, Thomas et al. 2017) while showing additional dependencies on light (**Fig. 1, f-h**). Understanding effects of abiotic conditions on the ZNGI is important, as this defines the fundamental niche of a species and provides a metric of its competitive ability (when ZNGIs are compared across species). Finally, (iv) the model also implies that TPCs are most skewed under nutrient replete conditions and optimal light levels and become more symmetric as resource limitation lowers growth rates.

We examined whether there is empirical support for these predictions about the effects of nutrients and light on *T*_*opt*_ (summarized in **Fig. 2a**). Regarding nutrients, we compiled data on individual TPCs measured at different phosphate levels for six species of phytoplankton (Thomas et al. 2017, Bestion et al. 2018) and estimated *T*_*opt*_ for each. For consistency with how these data were originally analyzed and to avoid circularity, we fit these TPCs using a Norberg TPC model (Norberg et al. 2001, Thomas et al. 2012) rather than the “double exponential” model (2) (Thomas et al. 2017). TPC fitting was conducted in R using the *growthTools* package v.1.2 (Kremer 2020). In four cases the estimated *T*_*opt*_ values for the Bestion et al. (2018) data were lower than their lowest experimental temperature (15°C); when this occurred, we replaced the *T*_*opt*_ estimate with 15°C (results were also qualitatively similar when these observations were simply excluded; not shown). Similarly, for light, we rely on previously compiled and analyzed trait data (Edwards et al. 2016), specifically *T*_*opt*_ values from Norberg TPC fits performed at different light levels and across 47 different species. To facilitate comparisons of qualitative patterns across species, which often differ in the position of their thermal niches, we then calculated relative *T*_*opt*_ values. For each species in each data set, this is the difference between its observed *T*_*opt*_ at particular light or phosphate levels and its mean *T*_*opt*_ across these abiotic conditions.

**Figure 2.**
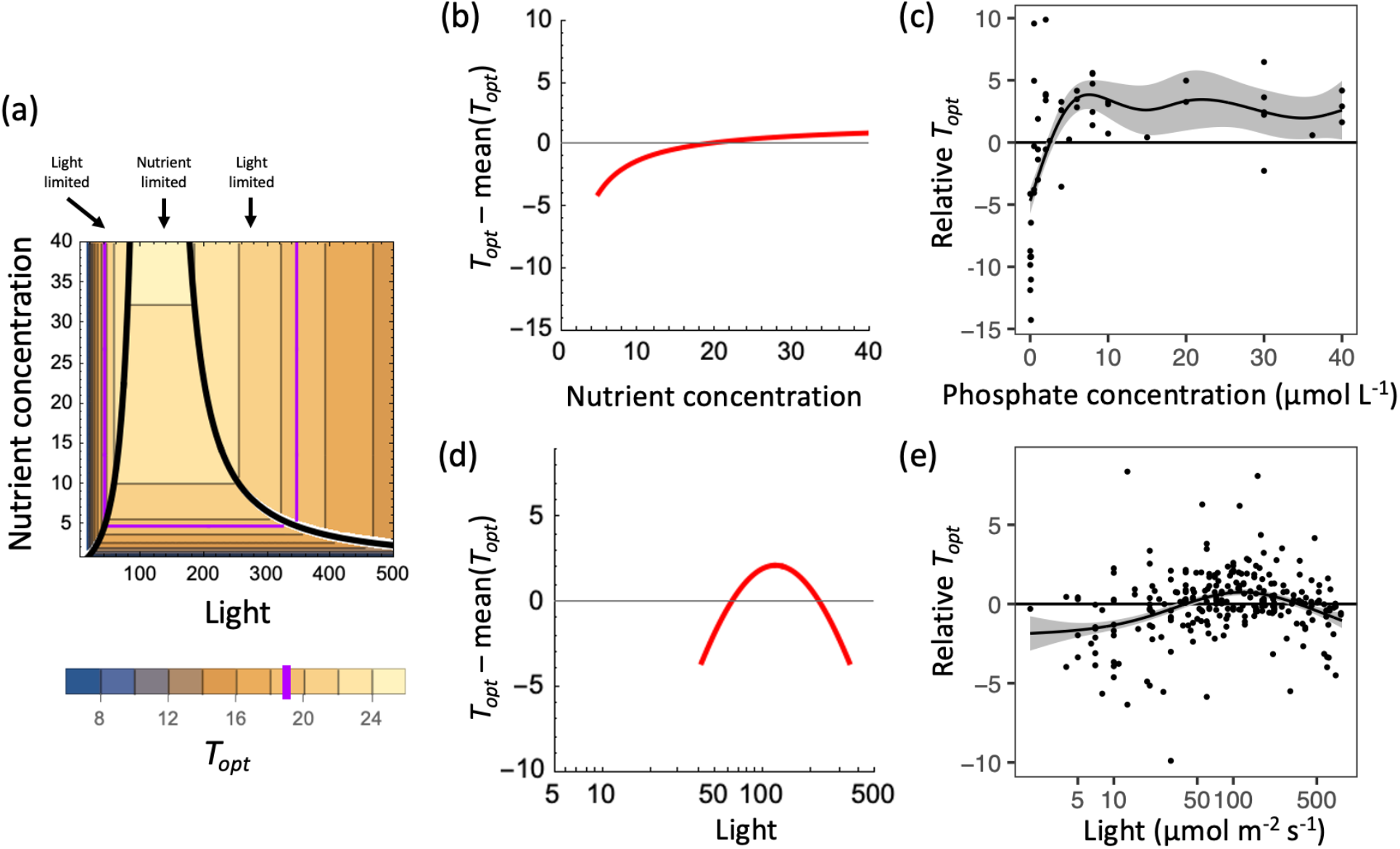
(a) A key prediction of our model is that *T*_*opt*_ varies with both nutrient and light supply, depending on which resource is limiting. Co-limitation by nutrients and light is indicated by the black solid line. Note that growth is only possible for combinations of nutrient and light yielding *T*_*opt*_ > 19°C (purple line). (b) *T*_*opt*_ is predicted to increase and saturate with increasing nutrient concentration (under optimal light – a vertical slice through (a)). This pattern is qualitatively consistent with an observed relationship across six different species (c) (Thomas et al. 2017, Bestion et al. 2018). (d) *T*_*opt*_ is also predicted to be a unimodal function of light level, peaking at *L*_*opt*_ (under replete nutrients – a horizontal slice through (a)). This is also qualitatively consistent with an analysis of 47 species (e) (Edwards et al. 2016). In (c, e), the y-axis displays relative *T*_*opt*_ (the difference between a species’ average *T*_*opt*_ across nutrient or light conditions and its observed *T*_*opt*_ at specific conditions). Trends shown by the black line and gray 95% confidence band in (c, e) arise from a generalized additive mixed model. Model parameters for (a, b, d), are given in Appendix S1: Table S1; curves in (b, d) are only plotted where conditions and traits lead to positive growth.

The qualitative prediction that *T*_*opt*_ is low at low nutrient levels and saturates at high nutrient levels (**Fig. 2b**) is consistent with the trends exhibited by empirical observations (**Fig. 2c**; Appendix S1: Table S2). We characterized these trends with a general additive mixed model or GAMM that included a random intercept for species, using the gamm4 package in R (version 0.2-6, (Wood and Scheipl 2020). Our second prediction, that *T*_*opt*_ peaks at intermediate light levels (**Fig. 2d**), is also qualitatively supported (**Fig. 2e**; Appendix S1: Table S3, same methods). To our knowledge, no sufficiently comprehensive experimental data exist for individual species to allow us to test our third and fourth predictions, relating the shape of the ZNGI to temperature, nutrients, and light, and the skewness of the TPC to nutrients and light.

### 2.2 Temperature-scaling of growth in photoautotrophs versus heterotrophs

Metabolic theory makes explicit predictions for the temperature scaling of the metabolic rate of photoautotrophs (activation energy *E* = 0.32 eV) and heterotrophs (*E* = 0.65 eV), based on differences in their rate-limiting biochemical processes (Allen et al. 2005). We hypothesize that these explanations for differences in temperature scaling between groups of organisms with distinct metabolic strategies may also apply to different temperature-dependent metabolic processes occurring within an organism that determine its overall growth. This leads to several specific predictions. First, photoautotrophs should exhibit birth and death rates that scale with temperature following MTE predictions for photosynthesis and respiration (respectively, 0.32 eV or 0.0498 for *b*_2_ and 0.65 eV or 0.101 for *d*_2_, invoking the approximation of Kremer et al. 2017). Second, the temperature sensitivity of birth in heterotrophs will be consistent with MTE values for respiration, and hence higher than in photoautotrophs. To test these predictions, and to explore thermal trait variation among species, we examined published datasets on the effects of temperature on growth rates of photoautotrophic (Thomas et al. 2016) and heterotrophic (Corkrey et al. 2016) microbes cultured under otherwise ideal conditions.

Prior to analysis, these data compilations required additional filtering and quality control (detailed in Appendix S3), to ensure that sample sizes were adequate and included enough temperatures to capture a unimodal response, to exclude thermophiles and taxa from Animalia and Fungi, to remove duplicates, and to omit data where factors other than temperature were limiting. We also removed cases where our model (2) fit poorly to the data (R^2^ < 0.85). After these measures, we were left with high quality fits of (2) to the thermal reaction norms of 248 microbial strains (113 heterotrophs, 110 autotrophs, and 25 mixotrophs). Curve fitting was performed using the growthTools package (Kremer 2020) in R (v. 4.3.3). This approach employs maximum likelihood methods and assumes normally distributed errors. Model fits provide estimates of the parameters of (2) for each microbial strain. All associated data and code are provided in the supplemental material.

Across microbial TPCs, temperature scaling parameters were quite variable (**Fig. 3a**). Generally, in photoautotrophs, *b*_2_ was lower than the predicted MTE value for the temperature sensitivity of photosynthesis, with a mean (95% CI) of 0.0345 (0.0283, 0.0420) (Appendix S1: Table S4). However, *d*_2_ was consistent with the predicted temperature sensitivity of respiration, 0.107 (0.088, 0.129) (Appendix S1: Table S4). In heterotrophs, *b*_2_ was also lower than predicted, 0.0629 (0.0539, 0.0733), although heterotrophs had a significantly higher *b*_2_ than the photoautotrophs (p < 0.0001), as predicted (Appendix S1: Table S4). If the generation of energy in heterotrophs arises via respiration, and *d*_2_ must be greater than *b*_2_ to produce a typical TPC, it is perhaps unsurprising that heterotroph parameter estimates are larger than those of autotrophs. Collectively, these results provide only partial support for our hypothesis. There was more variation in *b*_2_ and *d*_2_ than expected, which may relate to other factors ranging from the mundane (experimental noise) to issues deserving future study such as the evolutionary history or relatedness of strains, interactions with other environmental drivers, and cell size (Sal et al. 2015, Thomas et al. 2016, Kremer et al. 2017, Kontopoulos et al. 2020). Differences between photosynthesis and heterotrophic metabolism at least partially explain consistent differences in the TPCs of auto- and heterotrophs.

**Figure 3.**
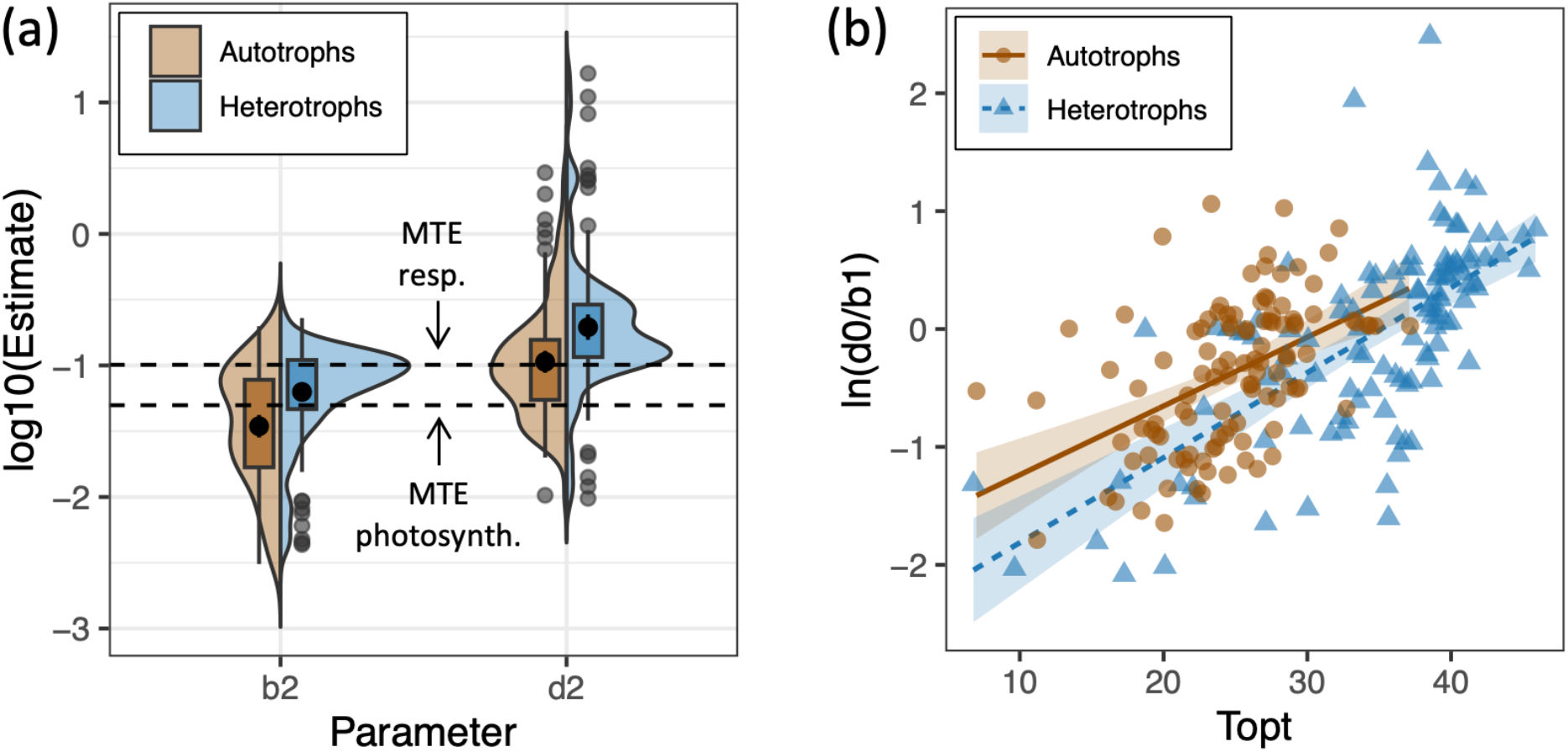
(a) Autotroph (brown) birth and death rates (*b*_2_ and *d*_2_) exhibit temperature sensitivities close to metabolic theory (MTE) predictions for photosynthesis and respiration (labeled dashed lines). These values are significantly higher in heterotrophs (blue), which have a thermal sensitivity of birth (*b*_2_) that approaches the MTE value for respiration (Appendix S1: Table S4). Distributions of trait estimates are shown both as box plots (including mean values, black points) with outliers (grey points) and as smoothed density functions. (b) A relationship between ln(*d*_0_/*b*_1_) and *T*_*opt*_ suggests a tradeoff constraining the shape of microbial thermal performance curves. Autotrophs and heterotrophs (brown vs. blue lines) have slightly different relationships (see Appendix S1: Table S5-S6).

### 2.3 Trade-offs constraining thermal performance

Understanding the parameter relationships that mathematically constrain the shape (height, breadth, skewness, etc.) of TPCs across species would help us draw valuable ecological and evolutionary inferences and predict responses to environmental warming. Two lines of evidence suggest that parameters in our model must be related to each other, which may reflect an underlying trade-off. First, consider (2) and assume that resources are not limiting and that populations differ only in their optimum temperature, 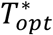. We show that under these conditions, any population with a higher 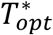 will grow faster at all temperatures than a population with a lower 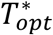 (Appendix S2: Section S4). This happens because in (2) increasing 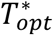 reduces death rates at all temperatures but does not affect *T*_*min*_. An important implication is that, without other constraints, selection on 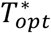 would lead to runaway evolution for ever higher values of 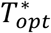 even in a constant environment (Appendix S2: Section S4), which is clearly not the case.

Second, an empirical analysis of the data from Section 2.2 shows that *d*_0_ and *b*_1_ — two of the parameters that determine *T*_*min*_(see Appendix S2: Section S2) — vary systematically with 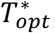, such that taxa with higher 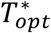 also tend to have higher *T* values, constraining niche widths. Specifically, within both photoautotrophs and heterotrophic microbes, we find that there is a strong log-linear relationship: In 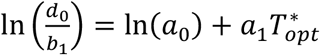, where In(*a*_0_) is the intercept and *a*_1_ the slope (**Fig. 3b**, Appendix S1: Table S5-S6). This relationship differs modestly between groups, with a slightly higher *a*_1_ in heterotrophs. These results are consistent with strong prior evidence in autotrophic microbes that 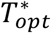 and *T*_*min*_ are related (Appendix 1: Fig. S1, Thomas et al. 2012, 2016, Kontopoulos et al. 2020). *T*_*min*_ increases more slowly with 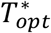 than *T*_*max*_, however, leading to a weak increase in niche width among taxa with higher 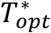 (Appendix 1: Fig. S1).

This relationship may capture an underlying physiological trade-off, or it may simply be an artifact resulting from a mismatch between the true cellular processes governing temperature-dependence and our mathematical description. In either case, uncovering and quantifying this relationship is important for two reasons. First, it suggests a possible change to (2) (replacing 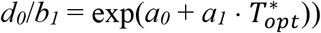 that captures a significantly, biologically meaningful constraint on variation in TPC shape, and removes the potential for runaway selection on 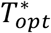 (Appendix S2: Section S2). Second, further consideration of this relationship highlights the importance of the relative sizes of *a*_1_ (slope of the relationship between ln 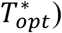 and *b*_*2*_ (strength of the temperature dependence of birth). If *a*_1_ > *b*_2_, *T*_*min*_ increases faster with 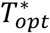 than *T*_*max*_, and we would expect species that specialize on high temperatures to have narrower thermal niches. In contrast, if *a*_1_ < *b*_2_, *T*_*min*_ increases more slowly than 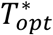, and we would expect species with higher 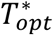 will also have wider (but finite) thermal niches (Appendix S2: Section S4). In either case, maximum growth rates would increase with increasing 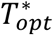, consistent with the ‘hotter is better’ hypothesis (Knies et al. 2009, Angilletta et al. 2010). However, only the first scenario (*a*_1_ > *b*_2_) would be consistent with long-standing hypotheses that propose a generalist-specialist tradeoff, where specialization on any particular temperature yields narrower thermal tolerance in addition to increased maximum growth (Angilletta 2009). Ultimately, we do not find evidence for this tradeoff, as the microbial data we analyze here suggest that *a*_1_ ≈ *b*_2_ for both trophic groups (Appendix S1: Table S7). This implies that niche width is unrelated, or at best weakly dependent, on 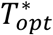, given the uncertainty in both parameter estimates (Appendix S1: Table S7). A direct analysis of trait estimates shows a weak increase in niche width with 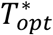 (Appendix S1: Figure S1b) (which would argue instead that *a*_1_ < *b*_2_), but this result is sensitive to sparse data at low 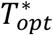. Despite this uncertainty, our analyses indicate how several fundamental hypotheses in thermal ecology can be connected within a single mathematical framework.

We have shown that the action of temperature and resource availability on population growth is more clearly understood by considering how these factors affect fundamental physiological processes of individual organisms (birth; death; photosynthesis; respiration). Next, we identify how properties of entire communities or ecosystems, especially scaling relationships characterizing their temperature- and resource-dependence, can be related to physiological constraints of individuals and populations.

## 3. Scaling from population to community-level relationships

Links between macroecological patterns and processes at lower levels of organization are scarce but important for understanding the fundamental rules of life (e.g., Wickman et al. 2024). Our model of temperature- and resource-dependent population growth suggests natural limits to the growth and production of entire communities as described by the geometrical envelope that bounds the collection of species-level TPCs. In this section, we establish an explicit connection between physiological constraints (temperature-dependence of photosynthesis) and these community-level patterns, including the Eppley curve (Eppley 1972, Bissinger et al. 2008, Kremer et al. 2017). Interactions between temperature and resource availability are also predicted to arise at this higher level of ecological organization. Finally, we show that these predictions are qualitatively consistent with field observations.

We can derive such a community-level relationship (or envelope) by manipulating our expression for the temperature- and resource-dependent growth rate of an individual species (**Box 1**, Appendix S2: Section S5). The result is an equation describing how the (maximum) net productivity of an entire community of species with different 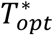 values will change across temperature and resource conditions, *μ*_*comm*_(*T, R*):

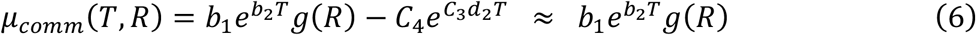

The first term of this equation, contributing positively to growth, is identical to (2), and depends on the temperature-dependent birth process limited by resource availability. The second part of this equation, a loss term, is again an exponential expression involving *d*_2_, but with some additional, temperature-independent constants involved (*C*_3_, *C*_4_, which are more lengthy algebraic expressions, see Appendix S2: Section S5). However, this loss term ends up being small and either independent or very weakly dependent on temperature under most realistic assumptions about parameter values and over the range of biologically relevant temperatures (Appendix S2: Section S5). The result is that when resources are not limiting, the community-level envelope *μ*_*comm*_(*T, R*) is nearly indistinguishable from an exponential function with a temperature scaling exponent of *b*_2_ (e.g., **Box 1**, Appendix S1: Fig. S4). As shown above (Section 2.2), typical values of *b*_2_ are consistent with the temperature-dependence of photosynthesis. This implies that we expect that the maximum growth rate of a group of phytoplankton that share other traits in common across temperature should follow a near exponential relationship with temperature corresponding to *b*_2_. This finding is consistent with analyses of empirical relationships (Kremer et al. 2017) and helps to explain the form of a fundamental relationship. Ultimately, this links physiological processes (birth and death, or photosynthesis and respiration) that underly population-level patterns (unimodal thermal performance curves) to known community-level relationships (the Eppley curve). This contrasts with previous approaches that assume this relationship *a priori* and impose it phenomenologically in defining species-level thermal performance curves (e.g., Norberg et al. 2001, Thomas et al. 2012).

In addition to proposing a more concrete basis for previously studied patterns, our results also offer novel predictions for how community-level responses like the Eppley curve are modified when resource levels are limiting. Ultimately, (6) provides an envelope surface that is a function of both temperature and limiting resource(s), which can include light and nutrients. For example, consider the effects of light limitation on the temperature-dependence of maximum interspecific growth rates in a community. Using parameter values characteristic of photoautotrophs arising from our preceding analyses, we can construct a hypothetical, but empirically grounded, envelope that depends on temperature and light (**Fig. 4a**). When light levels are optimal, the relationship between maximum growth rate and temperature is nearly exponential, exactly as described above (**Fig. 4b**). However, as light levels decline (*g*(*R*) decreases) the relationship weakens and eventually becomes nearly flat over biologically relevant temperatures (**Fig. 4c**). This result is qualitatively consistent with (Edwards et al. 2016), including their finding that specific carbon uptake (a proxy for growth) varied exponentially with temperature at high light levels (100-200 μmol m^-2^ s^-1^), but was unrelated to temperature at low light (20 μmol m^-2^ s^-1^) (reproduced in **Fig. 4 d,e**). As with light limitation, nutrient limitation can also cause a flattening of the scaling of community-level growth rates with temperature (**Fig. S2**).

**Figure 4.**
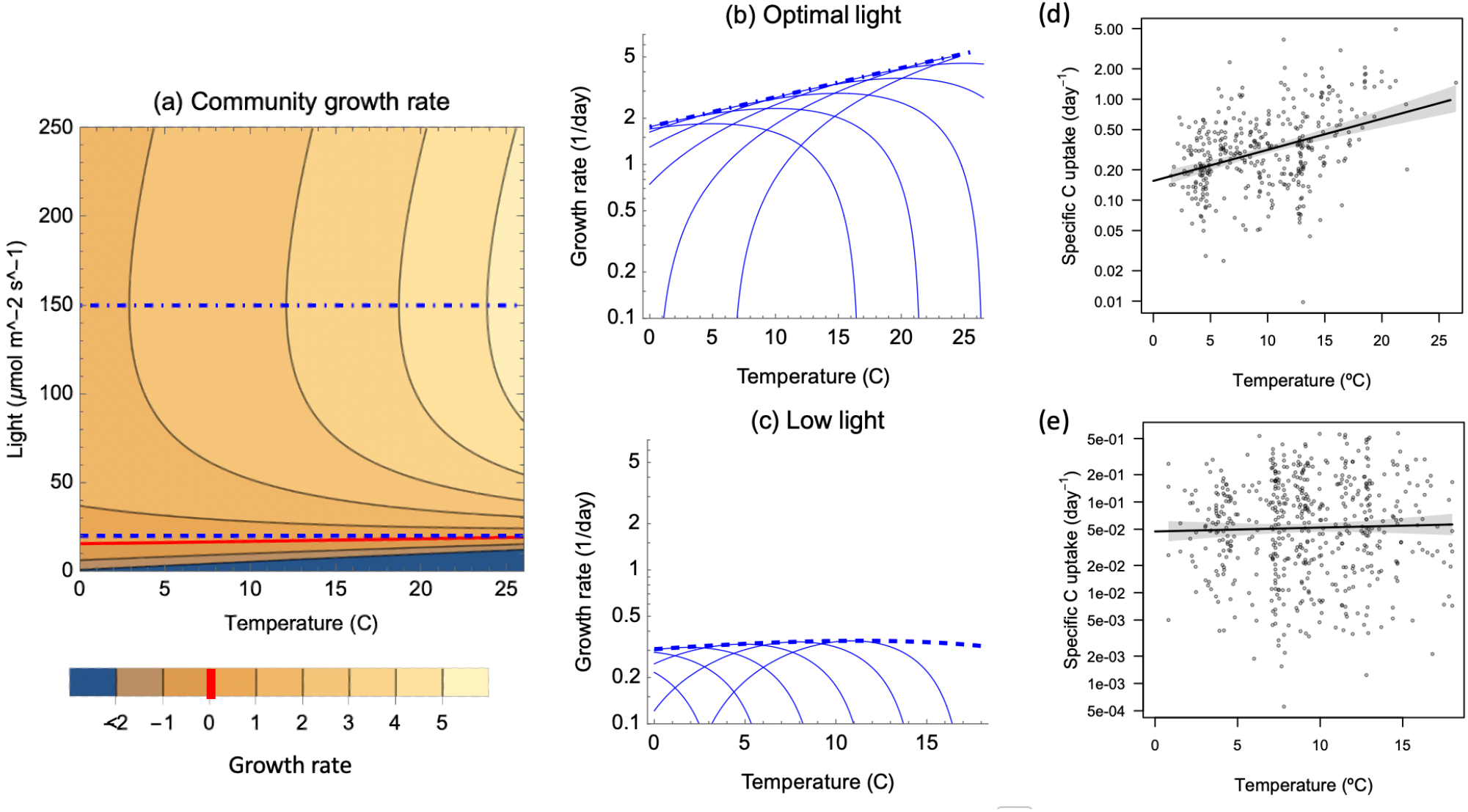
(a) The envelope function constraining the maximum interspecific growth rate of a community varies with both temperature and light, peaking at *L*_*opt*_ for any given temperature (assuming replete nutrients). As light approaches zero, growth rate becomes zero (red line) and eventually negative. Slices across this surface reveal the temperature sensitivity of growth rate at (b) optimal (dotted; 150 μmol m^-2^ s^-1^) and (c) low (dashed; 20 μmol m^-2^ s^-1^) and light levels. Example thermal performance curves of individual taxa with different 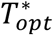 traits that contribute to these envelopes are drawn in thin lines. These patterns are qualitatively similar to the findings of (Edwards et al. 2016), demonstrating observational trends in the specific carbon uptake of marine phytoplankton communities at (d) high light (100-200 μmol m^-2^ s^-1^) and (e) low light (20 μmol m^-2^ s^-1^); republished with permission from the author. For (a)-(c), parameter values are representative of autotrophs, given earlier results: *b*_1_ = 3.97 (median across all photoautotrophs), *b*_2_ = 0.0498 (MTE prediction), *θ* = 0.065, *L*_*opt*_ = 150, *a*_0_ = 0.162 (Appendix S1: Table S5), *a*_1_ = 0.0584 (Fig. 3b autotrophs, Appendix S1: Table S5), *d*_2_ = *b*_2_ + 0.0563 (difference in median(*d*_2_) and median(*b*_2_) for photoautotrophs).

Although this qualitative agreement is exciting, several caveats are worth mentioning. First, the relationship between maximum growth rate and temperature at low light is not actually flat, but rather weakly unimodal in **Fig. 4c**: as temperatures continue to increase above 20°C, maximum growth rate declines more and more quickly, eventually becoming negative. Second, the result is somewhat sensitive to the particular choice of a “low” light level: lower or higher values produce relationships that are more strongly related to temperature. Finally, as in preceding results, the properties of this envelope surface depend on assumptions about key parameters, such as the value *b*_2_ and the difference between *a*_1_ and *b*_2_. Results in **Fig. 4** assume *b*_2_ is very slightly less than *a*_1_, consistent with empirical data and MTE predictions (see Appendix S1: Table S7). However, if this assumption is violated and *a*_1_ and *b*_2_ differ substantially, the relationship between maximum growth rate and temperature is unlikely to appear flat for any irradiance value over biologically relevant temperature ranges (**Appendix S1: Fig. S3**). Finally, this envelope analysis assumes that all community members share a common set of light traits; the presence of additional diversity in these traits would further influence the shape of the envelope function.

## Discussion

Forecasting the effects of global change on critical populations and communities requires linking abiotic conditions to their ecological consequences. This must be done in a way that balances the importance of capturing essential aspects of organismal physiology against the complexity and computational demands of scaling from individuals to populations and communities. Adopting a physiologically motivated approach, we propose population- and community-level models that capture the interactive effects of temperature and resources on ectothermic microorganisms. These models integrate ideas from multiple fields of research and produce several key predictions that are supported by our analysis of large compilations of empirical data on microorganisms. These results suggest intermediate-complexity scaling relationships that could increase the flexibility and accuracy of the biological components of global circulation models essential to predicting climate change effects. Our model and analyses provide several key theoretical connections and insights.

First, our model connects population growth rate to environmental conditions — a fundamental goal of ecology — for three of the most important environmental drivers of phytoplankton ecology (temperature, light, inorganic nutrients), incorporating non-trivial interactions between them. As none of these individual relationships is specific to phytoplankton (applying to all ectotherms, all photosynthetic organisms, and all life, respectively), it is likely that this model also applies to other taxa, as in the case of our previous work on temperature-nutrient interactions (Thomas et al. 2017). This is also consistent with the energetic principles underlying “metabolic meltdown”, where warming combined with food limitation further reduces organismal heat-tolerance (Huey and Kingsolver 2019). Quantifying these interactions allows us to define the abiotic niche of a species, which can then be used to mechanistically predict patterns of occurrence across real-world environments. This is, of course, possible only if the underlying relationships are adequately characterized via experimental work. Such experiments may be non-trivial, as the relationships are nonlinear and not well revealed by typical ANOVA designs without large numbers of treatment levels. Recent advances in experimental design offer some exciting solutions (Thomas and Ranjan 2024).

Second, our work emphasizes that critical phytoplankton traits that govern responses to specific abiotic factors (e.g., (Litchman and Klausmeier 2008)) depend unavoidably on suites of other abiotic factors, despite often being treated as idealized, independent quantities. For example, thermal performance curves under non-optimal nutrient or light conditions still have identifiable peaks (*T*_*opt*_ values), but the temperature at which these occur may be quite different from the 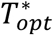 achieved under optimal conditions. These effects may emerge as a consequence of differential abiotic sensitivity of cellular processes, which are starting to be directly measured (e.g. metabolic rates, (Marañón et al. 2018)). Recognizing and accounting for these effects is critical both for making valid comparisons of traits across species, and for predicting the ecology of individual species in complex and dynamic abiotic environments. Similar to 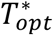, other canonical traits like half-saturation constants of nutrient-limited growth may be environment-dependent. For example, in our model nutrient limitation applies to the birth term only. The strength of this effect is governed by a temperature-independent half-saturation constant, determining the concentration of nutrients required for the population to achieve half of its maximum birth rate. In contrast, the resource concentration at which the population’s *net* growth rate is half of its maximum value may be quite different, and also will vary with temperature. This was implied, but not investigated in the model of (Thomas et al. 2017), and is increasingly a target of empirical investigation (Sunday et al. 2024). Other interactions may also exist: for example, our current model predicts that *L*_*opt*_ is independent of temperature or nutrient level. Comparative analyses suggest this is not the case (Fig. 2I, Edwards et al. 2016); the question deserves further study.

Third, we can address a long-standing debate in thermal ecology regarding constraints to thermal adaptation. Although we find that maximum growth rates increase in species adapted to warmer temperatures (with higher 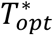), such species do not have consistently narrower thermal niches than cold-adapted species, and may in fact have wider niches on average (Knies et al. 2009, Thomas et al. 2016, Kontopoulos et al. 2020). Thus, we find no evidence of a clear generalist-specialist tradeoff. The formulation of our model does reveal a strong relationship between 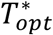 and the ratio between temperature-independent loss (*d*_0_) and birth rates at 0°C (*b*_1_). Such a relationship could be indicative of an underlying tradeoff, or simply a mathematical property arising from the particular formulation and parameterization of our model (2), possibilities that warrant further investigation. Either way, these constraints are empirically quantifiable and important, because they can influence the properties of envelope functions bounding community-level performance.

Lastly, our model is consistent with the idea that the temperature-dependence of performance at the organismal level can be understood by considering the combined effects of essential sub-organismal processes, which may have distinct temperature and nutrient sensitivities (Thomas et al. 2017, Marañón et al. 2018). This is inherent in predictions from metabolic theory that autotrophs and heterotrophs have different rate-limiting processes for obtaining energy, and that the temperature dependence of these rates explains the different metabolic scaling of these two groups (Allen et al. 2005, Kremer et al. 2017). Our results agree: the temperature sensitivities of birth and death in photoautotrophs are consistent, respectively, with the temperature scaling of photosynthesis and respiration from metabolic theory, although substantial species-specific variation remains. We also show that this result is not a given, as an otherwise very similar group of organisms (heterotrophic microbes) show distinctly different temperature sensitivities, including a temperature sensitivity for birth that is more closely aligned with respiration. This suggests that, overall, our model can be used to understand temperature-dependence for both groups of organisms, but that the underlying processes giving rise to particular growth responses likely have a different mechanistic basis and temperature dependence. An interesting extension of this work would be to study mixotrophs, which engage in a mix of heterotrophy and photosynthesis in amounts that vary under different conditions (Wilken et al. 2013, Wieczynski et al. 2023). Such an investigation might even more cleanly reveal how different patterns of thermal sensitivity arise under different feeding modes.

Despite this success, there are also numerous ways to expand or adapt the models we present and further integrate physiology and ecology. Some promising avenues for future improvements include:

i. Temperature-dependent population birth rates are probably not truly exponential. Over a wide enough range of temperatures, even birth rates must be unimodal. Alternatives used in the theoretical literature include Gaussian and logistic temperature-dependencies (e.g. Amarasekare and Savage 2012, Bieg and Vasseur 2024). An alternate modeling approach based on energetics also generates TPCs that change with food availability and uses a Gaussian energy acquisition function (Huey and Kingsolver 2019). These alternatives could be explored in an analysis similar to ours. However, as long as the intersection between the birth and death curves occurs in the range of temperatures where the birth curve is approximately exponential (e.g. the ascending portion of a Gaussian curve or a logistic curve before its inflection point), our simpler model is likely to remain a reasonable approximation. Historically, it has been difficult to experimentally distinguish birth and death rates in microorganisms like phytoplankton, so empirical results on the temperature dependence of birth and death in phytoplankton are extremely limited. Newly developed methods are overcoming this challenge and promise insights into the actual functional form of the birth and death curves (Baker and Geider 2021).
ii. Interactions between multiple resources (including light and nutrients) are more complex than Liebig’s law of the minimum implies, and often organisms may be co-limited by two or more resources. For example, the construction and operation of light harvesting machinery requires resources. These issues have been previously highlighted and some alternatives have been suggested (Klausmeier et al. 2004, Schade et al. 2005, Sperfeld et al. 2012, Bonachela et al. 2016). However, these changes matter most where there is a transition between limiting factors; if this occurs across a relatively narrow range of environmental space the practical consequences may not be large.
iii. Growth is governed by multiple physiological processes including resource uptake, assimilation and storage. The commonly used Droop model improves on the older Monod model by making phytoplankton growth arise from uptake rates and intracellular storage (quotas) (Droop 1972). It would be interesting to consider how temperature-nutrient-light interactions affect such a model, and whether additional allocation trade-offs are revealed (Ward 2017). For example, (Bieg and Vasseur 2024) consider effects of temperature on uptake and assimilation, and how these in turn affect the temperature dependence of per capita growth rate and carrying capacity.

All three of these avenues for model improvements reflect areas where added complexity may increase model accuracy over portions of parameter or environmental space. However, there is also considerable value in model simplicity, so we consider these to be opportunities rather than major drawbacks to the current formulation. Overall, our goal remains to obtain reasonable approximations of the collective behavior and ecology of groups of organisms as environmental conditions vary in space and time, rather than to derive models that resolve in great detail the specific physiology of many individuals comprising populations and communities.

## Conclusion

Advancing ecology, including both basic understanding and our ability to make reliable predictions of the state and functioning of future ecological systems, requires developing scalable, mechanistic models that account for interactive abiotic effects. Such models should ideally be physiologically motivated and able to capture critical empirical patterns, yet also parsimonious. We have developed an example of such a model, focusing on the interactive effects of temperature and resources on population- and community-level patterns and processes. We derived a series of predictions emerging from this model at both scales and demonstrated that these predictions are consistent with data on the growth and production of diverse groups of microorganisms, as well as predictions from other bodies of theory. The model and its results are likely to be generalizable to other ectotherms, perhaps with further adjustments to capture essential features or unique physiological details such as population structure or consumer-resource dynamics. Finally, we argue that this model could be incorporated into large scale ecosystem models critical to predicting the effects of ongoing environmental change, including warming and altered resource distributions, on microbial processes and biogeochemistry.

## Supporting information

Appendix 2: Mathematical analyses

Appendix 3: Data filtering and quality control

Appendix 4: Guide to code and supporting analyses

## Acknowledgments

We thank Kyle F. Edwards for advice and suggestions related to interpreting temperature-light interactions and field data, and financial support from NSF grant OCE-1638958 to EL and CAK. This is W. K. Kellogg Biological Station contribution number XXX.

## Author Contributions

CTK and MKT conceived of the paper; CTK and CAK performed the mathematical analyses; CTK conducted all statistical analyses; CTK and MKT drafted the manuscript; all authors contributed intellectually and to revising the manuscript.

## Box 1.

**Community envelope derivation and assumptions**

Our goal is to derive an expression for the envelope bounding the maximum growth rate achievable by the most productive (fastest growing) member of a community under any given temperature and resource environment. We will denote this relationship as *μ*_*comm*_(*T,R*) and obtain it by analyzing our existing expression for the temperature- and resource-dependent growth rate of an individual species, *μ*(*T,R*), given by (2). Full mathematical details are provided in Appendix S2: Section S5, with accompanying visuals in Appendix S1: Fig. S4. Note that *μ*(*T,R*) for any given species depends on its traits, e.g., 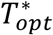, *b*’s, *d*’s, and parameters of its functional responses, such as the half-saturation constant *K*.

Accomplishing this task requires several key mathematical assumptions, consistent with ideas used to discuss the empirical pattern motivating this analysis, the Eppley curve (Eppley 1972, Moisan et al. 2002, Kremer et al. 2017). First, we assume that maximum growth rate of an entire community at any moment in time can be described by the highest-performing species in the current environment (e.g., the one that grows faster than any other species). Second, we assume that as environmental conditions change, the identity of this optimal species also changes immediately. This implies that adequate functional diversity exists in an accessible species pool (such that the theoretically ‘optimal’ species exists or can rapidly evolve), and that community dynamics occur faster than rates of environmental change (such that the ‘optimal’ species dominates the community at all times). Then, by deriving relationships that characterize how the properties of ‘optimal’ species change across environmental gradients, we can obtain the desired community-level scaling relationship.

Now, we seek to determine how the (maximum) productivity of an entire community varies across both temperature and resource conditions; we denote this relationship as *μ*_*comm*_(*T, R*). To obtain this expression, we assume that the species pool contains a set of species that share common values for all parameters in (2) <u>except</u> their 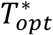 traits, which vary continuously across species. We also assume that the *d*_0_/*b*_1_ vs. 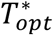 tradeoff that we found in Section 2.3 holds. Then, using calculus (Appendix S2: Section S5), we obtain equation (6), which determines envelope show by the red line in Appendix S1: Fig. S4.

