## Appendix 1: Supplemental Tables and Figures for "How interactions between temperature and resources scale from populations to communities"

### Appendix S1: Supplemental figures & tables

**Table S1.** Model parameters used in Fig. 1, and Fig. 2, a, b, and d. See code in Appendix S4.

| Variable | Value |
|---|---|
| $d_0$ | 0.1 |
| $d_1$ | 0.01 |
| $d_2$ | 0.11 |
| $b_2$ | 0.05 |
| $T_{opt}$ | 25 |
| $k$ | 2 |
| $\mu_{max}$ | 1.2 |
| $\alpha$ | 0.03 |
| $L_{opt}$ | 120 |

**Table S2.** Summary of GAMM results for Relative $T_{opt}$ ~ Phosphate (Fig. 2c); scale estimate = 18.937, n = 52, adj-$R^2$ = 0.384.

Fixed Effects

| | Estimate | Std. error | t value | p-value |
|---|---|---|---|---|
| Intercept | -3e-15 | 0.6035 | 0 | 1.00 |
| | edf | Ref.df | F value | p-value |
| s(Phosphate) | 3.925 | 3.925 | 7.374 | < 0.0001 |

Random Effects

| | n groups | Variance | Std. dev |
|---|---|---|---|
| Intercept \| Species | 6 | 0.00 | 0.000 |
| s(Phosphate) | 4 | 1144.18 | 33.826 |
| Residual | | 18.94 | 4.352 |

**Table S3.** Summary of GAMM results for Relative $T_{opt}$ ~ Light (Fig. 2e); scale estimate = 3.8557, n = 266, adj-$R^2$ = 0.122.

Fixed Effects

| | Estimate | Std. error | t value | p-value |
|---|---|---|---|---|
| Intercept | -1.7e-16 | 0.1204 | 0 | 1.00 |
| | edf | Ref.df | F value | p-value |
| s(log10(irradiance)) | 3.454 | 3.454 | 11.2 | < 0.0001 |

Random Effects

| | n groups | Variance | Std. dev |
|---|---|---|---|
| Intercept \| Species | 47 | 0.00 | 0.000 |
| s(log10(irradiance)) | 4 | 25.328 | 5.033 |
| Residual | | 3.856 | 1.964 |

**Table S4.** Estimates of the mean temperature scaling of birth ($b_2$) and death ($d_2$) rates from the fit of E.2 to a database of autotrophic (A) and heterotrophic (H) microbial thermal performance curves, comparisons of these values to predictions from metabolic theory (MTE), and associated hypothesis tests.

**Hypothesis tests**

| Parameter | Estimate (95% CI) | Prediction ($b_{2,MTE}$) | $H_0$: ($b_2$ - $b_{2,MTE}$) = 0 | | $H_0$: $b_{2,A}$ - $b_{2,H}$ = 0 | |
|---|---|---|---|---|---|---|
| | | | t | p-value | F | p-value |
| $b_{2,A}$ | 0.0345 (0.0283, 0.0420) | 0.0498 | -3.689 | 0.00035 | 22.74 (1, 221) | < 0.0001 |
| $b_{2,H}$ | 0.0629 (0.0539, 0.0733) | 0.101 | -6.134 | < 0.0001 | | |

| Parameter | Estimate (95% CI) | Prediction ($d_{2,MTE}$) | $H_0$: ($d_2$ - $d_{2,MTE}$) = 0 | | $H_0$: $d_{2,A}$ - $d_{2,H}$ = 0 | |
|---|---|---|---|---|---|---|
| | | | t | p-value | F | p-value |
| $d_{2,A}$ | 0.107 (0.088, 0.129) | 0.101 | 0.535 | 0.594 | 16.66 (1, 221) | < 0.0001 |
| $d_{2,H}$ | 0.195 (0.157, 0.244) | - | - | | | |

**Table S5.** Coefficient estimates and confidence intervals for $\ln(d_0/b_1) \sim a_0 + a_1 \cdot T_{opt}$ for photoautotrophs. $R^2 = 0.23$, F-value = 32.86 (1, 108).

| Coefficient | Estimate (95% CI) | Std. Error | t-value | p-value |
|---|---|---|---|---|
| $\ln(a_0)$ | -1.820 (-2.33, -1.31) | 0.256 | -7.112 | < 0.0001 |
| $a_1$, effect of $T_{opt}$ | 0.0584 (0.0382, 0.0785) | 0.010 | 5.732 | < 0.0001 |

**Table S6.** Coefficient estimates and confidence intervals for $\ln(d_0/b_1) \sim a_0 + a_1 \cdot T_{opt}$ for heterotrophs. $R^2 = 0.43$, F-value = 84.22 (1, 111).

| Coefficient | Estimate (95% CI) | Std. Error | t-value | p-value |
|---|---|---|---|---|
| $\ln(a_0)$ | -2.534 (-3.08, -1.98) | 0.277 | -9.135 | < 0.0001 |
| $a_1$, effect of $T_{opt}$ | 0.0721 (0.0566, 0.0877) | 0.008 | 9.177 | < 0.0001 |

**Table S7.** Comparing the estimate and confidence interval of the $T_{opt}$ effect (parameter $a_1$) with average $b_2$ values (and confidence intervals) taken across species/TPCs within auto- and heterotrophs.

| | n | Mean $b_2$ (95% CI) | $a_1$ (95% CI) |
|---|---|---|---|
| Photoautotrophs | 110 | 0.0345 (0.0283, 0.0420) | 0.0584 (0.0382, 0.0785) |
| Heterotrophs | 113 | 0.0629 (0.0539, 0.0733) | 0.0721 (0.0566, 0.0877) |

(a)

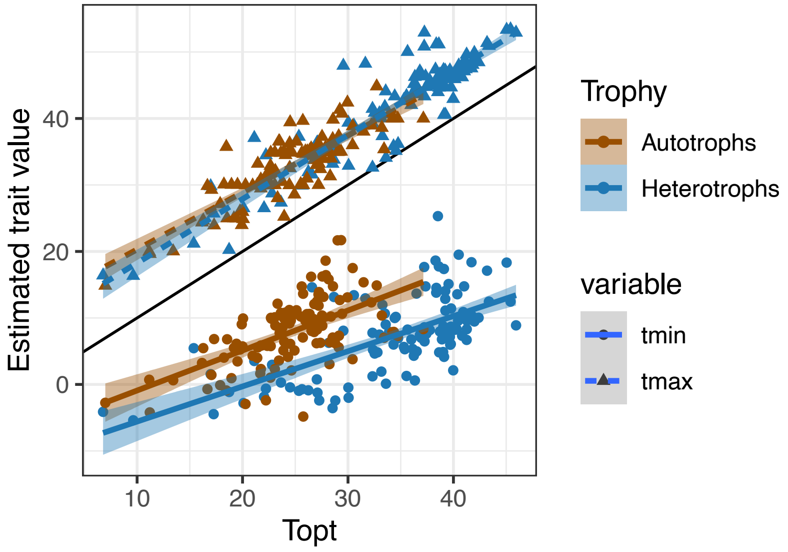


(b)

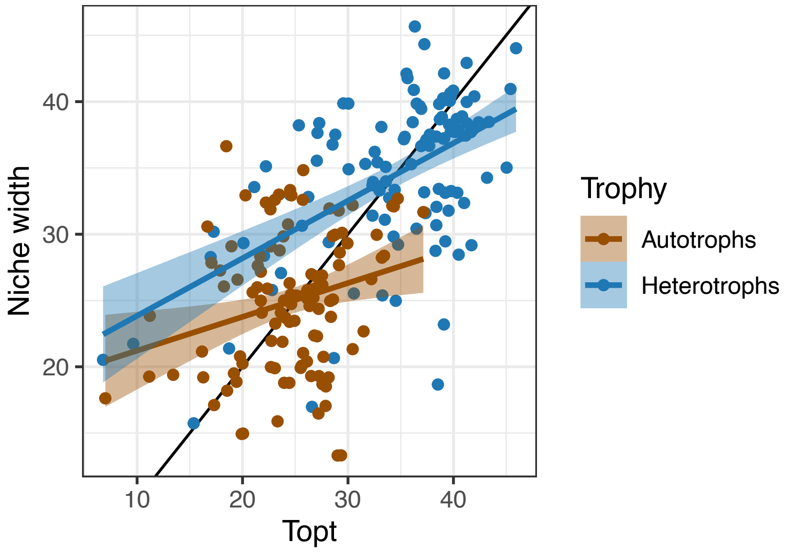


**Figure S1**. (a) $T_{max}$ increases with optimum temperatures ($T_{opt}$) such that it is consistently 7-8°C higher than $T_{opt}$. $T_{min}$ increases more slowly with increasing $T_{opt}$. The black solid line shows a 1:1 relationship; above this line are the $T_{max}$ values (triangles; dashed lines) and below it are estimates of $T_{min}$ (dots, solid lines). Trends are displayed separately for photoautotrophs (brown) and heterotrophs (blue). In (b) we see that the responses of $T_{min}$ and $T_{max}$ combine to produce a weak overall trend of increasing niche with for curves with higher $T_{opt}$. However, the strength of this relationship is influenced substantially by the relatively sparse data for $T_{opt} \leq 15$°C.

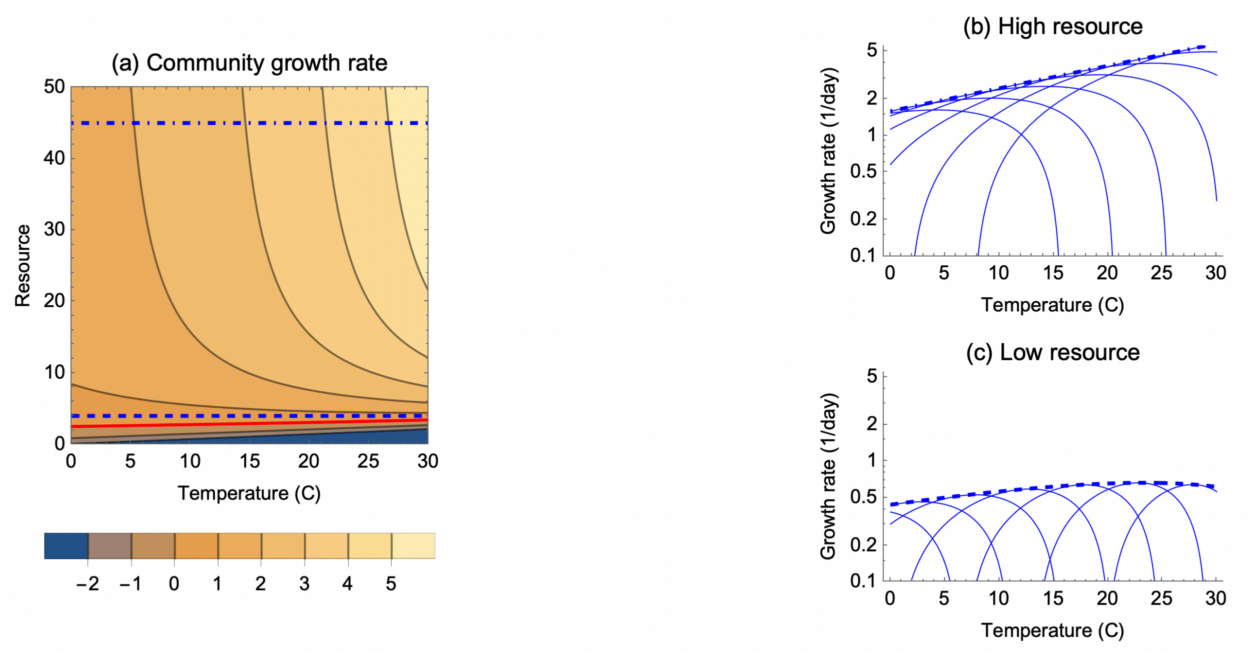


**Figure S2.** (a) The envelope function constraining the maximum interspecific growth rate of a community varies with both temperature and resource, similar to Fig. 4. As resource approaches zero, growth rate becomes zero (red line) and eventually negative. Slices across this surface at (b) optimal (dotted; $R = 45$) and (c) low (dashed; $R = 4$) resource levels reveal the temperature sensitivity of growth rate. Performance curves of taxa with different $T^*_{opt}$ traits underlying these envelopes are drawn in thin lines. Parameter values are as in Fig. 4.

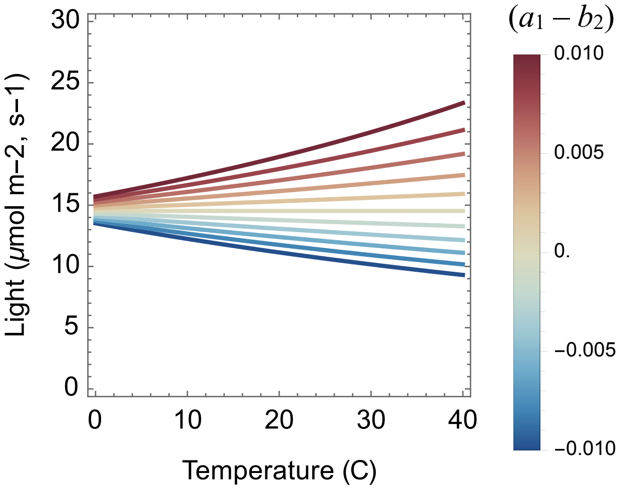


**Figure S3.** The community-level zero net growth isocline (ZNGI) as a function of irradiance and temperature depends on the difference between $a_1$ and $b_2$. Individual lines represent the community ZNGI for a different value of the difference between $(a_1 - b_2)$, indicated by color. In each scenario, growth rate would be positive above a given curve and negative below. Note that when $a_1 = b_2$, the ZNGI is flat (independent of temperature). When $a_1 > b_2$ (red), the lowest irradiance that still allows growth is higher at higher temperatures. When $a_1 < b_2$, the lowest irradiance permitting growth actually declines at higher temperatures. For the overall relationship between maximum growth rate and temperature to appear effectively flat at low irradiance levels, it is necessary that $a_1 > b_2$ by a fairly small amount. This is plausible for autotrophs (Fig. S1, Table S7), although uncertainty remains. Parameters are as in Fig. 4.

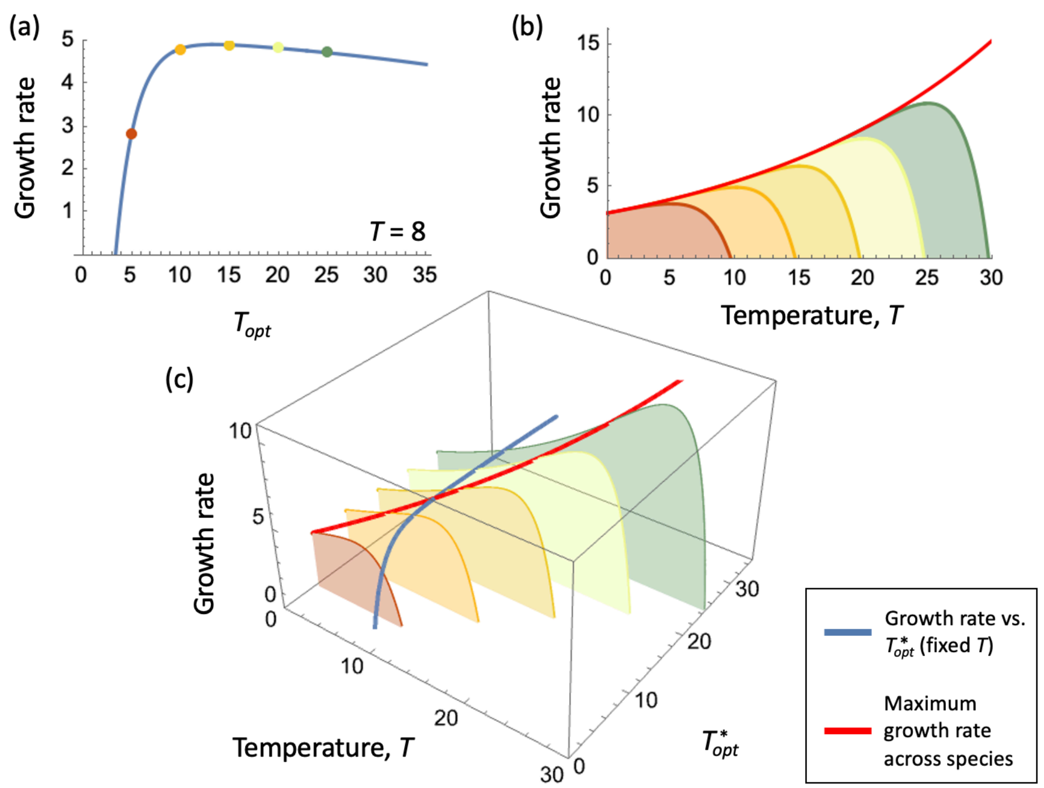


**Figure S4**. (a) Illustrates how growth rate (blue line) at fixed temperature $T$ varies with $T^*_{opt}$; colored dots correspond to the individual, example thermal performance curves, TPCs, drawn in (c). (b) Shows how the envelope function for maximum growth rate across species (red line) bounds the entire family of TPCs across a range of temperatures. (c) 3D graphic demonstrating how the maximum growth rate (across $T^*_{opt}$, for fixed $T$, blue line) leads to the envelope function (red line).
