## Appendix 2: Mathematical analyses for "How interactions between temperature and resources scale from populations to communities"

**Appendix S2: Mathematical analysis of the double exponential model and its extensions**

Section S1: Basic model (no resource limitation)

As defined in Thomas et al. 2017, the double exponential model attributes change in net population growth rate as a function of temperature, $\mu(T)$, to the difference between birth and death processes, following

$$\mu(T) = b_1 e^{b_2 T} - (d_0 + d_1 e^{d_2 T}) \qquad (S1)$$

(This is identical to (1) in the main text, under the assumption that resources are not limiting; i.e. that $g(R) = 1$). Here, the birth term depends on $b_1$ (birth rates at $T = 0$ °C) and $b_2$ (determining the exponential increase in birth rates with temperature, $T$). The death term consists of two components: first, a temperature-independent mortality rate, $d_0$, and second, a temperature-dependent process that depends on $d_1$ (death rate at $T = 0$ °C) and $d_2$ (setting the exponential increase in death rate with temperature). All parameters are restricted to be greater than zero. Additionally, for (Equation S1) to produce canonical unimodal thermal reaction norms, $b_2 < d_2$. This model is similar in form to a model proposed by (Logan et al. 1976), with the addition of the $d_0$ term. Next, we demonstrate how to calculate or approximate key properties of this thermal performance curve (TPC) equation.

Section S2: Cardinal thermal traits

Frequently, it is useful to determine the optimal temperature ($T_{opt}$) at which a species achieves its highest growth rate ($\mu_{max}$), as well as the temperatures at which it ceases to be able to grow ($T_{min}$ and $T_{max}$), and the distance between them (niche width). Here we consider the calculation of these cardinal thermal traits from the basic double exponential model (Equation S1).

*S2.1 Optimum temperature, $T_{opt}$*

First, it is possible to reparametrize (Equation S1) to depend explicitly on $T_{opt}$ (see (O'Donnell et al. 2018)). This involves calculating the temperature at which it achieves its maximum value, or $\mu_{max}$, by taking the derivative of (Equation S1), setting it equal to zero, and solving for $T$

$$T^*_{opt} = \frac{1}{d_2 - b_2} \ln\left(\frac{b_1 b_2}{d_1 d_2}\right) \tag{S2}$$

(We note that, as in the main text, we indicate that this optimum temperature is defined assuming no resource limitation by using the '*' symbol). Having obtained this analytical representation for $T^*_{opt}$, we can then select a parameter of Equation S1, in this case $d_1$, and express that parameter as a function of $T^*_{opt}$. Substituting the resulting expression into Equation S1, we obtain

$$\mu(T) = b_1 e^{b_2 T} - \left(d_0 + \frac{b_1 b_2}{d_2} e^{(b_2 - d_2)T^*_{opt}} e^{d_2 T}\right) \tag{S3}$$

where the first term treats birth as an increasing exponential function of temperature.

*S2.2 Maximum growth rate, $\mu_{max}$*

This value is obtained simply by evaluating Equation S3 at $T = T^*_{opt}$, yielding:

$$\mu_{max} = \frac{b_1(d_2 - b_2)}{d_2} e^{b_2 T^*_{opt}} - d_0 \tag{S4}$$

We could use this result to further reparameterize Equation S3, to obtain an equation for the double exponential model TPC that includes an explicit parameter for maximum growth rate. This involves solving Equation S4 for $b_1$, then substituting the result into Equation S3, yielding

$$\mu(T) = \left(\frac{d_2}{d_2 - b_2} e^{b_2(T - T^*_{opt})} - \frac{b_2}{d_2 - b_2} e^{d_2(T - T^*_{opt})}\right)(\mu^*_{max} + d_0) - d_0 \tag{S5}$$

We note here that $T^*_{opt}$ is only the optimum temperature under optimal conditions (no resource limitation), and $\mu^*_{max}$ is specifically the maximum growth rate without resource limitation and where $T = T^*_{opt}$.

*S2.3 Minimum temperature, $T_{min}$*

Unfortunately, it does not appear possible to obtain exact analytical expressions for $T_{min}$. However, a reasonable analytical approximation exists. As $T$ declines far below $T^*_{opt}$, the term $e^{d_2T}$ will approach zero faster than $e^{b_2T}$, because $d_2 > b_2$. So, if we assume that $e^{d_2T} \approx 0$, Equation S3 becomes:

$$\mu(T) \approx b_1 e^{b_2T} - d_0 \quad \text{(S6)}$$

Setting Equation S6 equal to zero and solving for $T$, we obtain an analytical approximation for $T_{min}$:

$$T_{min} \approx \frac{ln(d_0/b_1)}{b_2} \quad \text{(S7)}$$

This approximation will be reasonably good as long as $T_{min}$ is not too close to $T^*_{opt}$. Generally speaking, this approximation will yield $T_{min}$ values that are lower than the true $T_{min}$, which can be calculated using numerical methods.

*S2.4 Maximum temperature, $T_{max}$*

Similarly, it does not appear possible to obtain exact analytical expressions for $T_{max}$. However, as $T$ increases to large values, the exponential terms $e^{d_2T}$ and $e^{b_2T}$ should become significantly larger than $d_0$. If we thus assume that $d_0 = 0$, we can solve Equation S3 for the value of $T$ where growth rate falls to zero, obtaining:

$$T_{max} \approx T^*_{opt} + \frac{\ln(d_2/b_2)}{d_2 - b_2} \quad \text{(S8)}$$

This approximate $T_{max}$ value should generally be higher than the true $T_{max}$ value, which can again be calculated using numerical methods.

*S2.5 Niche width, ω*

Niche width can be approximated by taking the difference between the $T_{max}$ and $T_{min}$ approximations,

$$\omega \approx \left(T_{opt}^{*} + \frac{\ln(d_2/b_2)}{d_2 - b_2}\right) - \frac{ln(d_0/b_1)}{b_2} \quad \text{(S9)}$$

This niche width approximation will be wider than the true niche width of a species, given the biases of the underlying $T_{min}$ and $T_{max}$ approximations.

Section S3: Mathematical notes on light limitation

The effect of light $L$ (also termed irradiance) on growth rate is often captured using the unimodal Eilers-Peeters equation (Eilers and Peeters 1988):

$$\mu(L) = \frac{\mu_{max}\, L}{\frac{\mu_{max}}{\alpha\, L_{opt}^2} L^2 + \left(1 - 2\frac{\mu_{max}}{\alpha\, L_{opt}}\right) L + \frac{\mu_{max}}{\alpha}} = \mu_{max} \cdot f_L(L) \quad \text{(S10)}$$

where $L_{opt}$ is the light level where growth rates peak, $\alpha$ is the slope of the relationship between growth rate and light when light is near zero, and $\mu_{\max}$ is the maximum growth rate. Rather than letting net population growth rate be limited by light, we wish to consider how light affects birth rates alone. We next consider how to incorporate this standard relationship into our birth-death model, which is (2) in the main text.

In the case of nutrient limitation, we used the Monod equation, which consists of the product of a parameter $b_{max}$ specifying the maximum birth rate and $f_R = R/(R+K)$. It was thus easy to define the maximum birth rate as $b_{max} = b_1 e^{b_2 T}$, and then to observe that the remaining term $f_R$ is a temperature-independent, saturating function of resource $R$ that asymptotes to 1 in the limit of high resource levels.

In the case of light limitation and the Eilers-Peeters relationship, the situation is more complicated, because the maximum rate term, $\mu_{max}$, appears in both the numerator and denominator of (Equation S10). We can factor the maximum rate term out of the numerator and replace it with $b_{max} = b_1 e^{b_2 T}$, as above. However, what do we do about $\mu_{max}$ in the denominator? If we similarly substitute $b_1 e^{b_2 T}$ into the denominator, then we have

$$b(T, L) = b_1 e^{b_2 T} \cdot \frac{L}{\frac{b_1 e^{b_2 T} \cdot}{\alpha\, L_{opt}^2} L^2 + \left(1 - 2\frac{b_1 e^{b_2 T} \cdot}{\alpha\, L_{opt}}\right) L + \frac{b_1 e^{b_2 T} \cdot}{\alpha}} = b_{max} \cdot f_L(T, L) \qquad \text{(S11)}$$

In this formulation, $f_L$ is not independent of temperature, which makes it more difficult to separate out the effects of temperature and resource limitation on population growth. However,

$$\frac{\partial}{\partial L}\left[b_{max} \cdot f_L(T, L)\right]\Big|_{L=0} = \alpha,$$

implying that the parameter $\alpha$ retains its interpretation in Equation S10 as the slope of the relationship between birth rate and light level at near-zero light.

Alternatively, we could treat the $\mu_{max}$ values in the numerator and denominator of Equation S10 as two independent parameters, $b_{max}$ and $b'$, respectively. If we let $b_{max} = b_1 e^{b_2 T}$ as above, but assume that $b'$ is independent of temperature, and define a new parameter $\theta = b'/\alpha$, the resulting relationship is

$$b(T, L) = b_1 e^{b_2 T} \cdot \frac{L}{\left(\frac{1}{\theta \cdot L_{opt}^2}\right) L^2 + \left(1 - \frac{2}{\theta \cdot L_{opt}}\right) L + \left(\frac{1}{\theta}\right)} = b_{max} \cdot f_L(L) \qquad \text{(S12)}$$

Although this is no longer equivalent to Equation S11, the resulting $f_L$ has two useful features: (i) it is independent of temperature, and (ii) it is still a unimodal function of light, peaking at $L_{opt}$. This is the formulation used in the main text. However, one drawback is that we cannot interpret

$\theta$ as easily as $\alpha$ in Equation S11. Inspecting $\frac{\partial}{\partial I}\left[b_{max} \cdot f_L(L)\right]\Big|_{L=0}$ using Equation S12, we find that

$$\frac{\partial}{\partial L}\left[b_1 e^{b_2 T} \cdot \frac{L}{\left(\frac{1}{\theta \cdot L_{opt}^2}\right)L^2 + \left(1 - \frac{2}{\theta \cdot L_{opt}}\right)L + \left(\frac{1}{\theta}\right)}\right]\Bigg|_{L=0} = \theta \cdot b_1 e^{b_2 T}$$

Consequently, the slope of the relationship between birth rate and light at low light levels is no longer a constant, but rather an exponential function of temperature.

Section S4: Potential constraints on model parameters

*S4.1 Motivation – demonstrating runaway selection on $T_{opt}$*

Consider the relationship between two species, *A* and *B*, that differ only in their optimum temperature traits, $T_{opt,A}^*$ and $T_{opt,B}^*$, where $T_{opt,B}^* > T_{opt,A}^*$. In particular, examine the relationship between the growth rates of these species across a range of temperatures, $\mu_A(T)$ and $\mu_B(T)$.

$$\mu_B(T) - \mu_A(T) = \left(b_1 e^{b_2 T} - \left(d_0 + \frac{b_1 b_2}{d_2} e^{(b_2 - d_2)T_{opt,B}^*} e^{d_2 T}\right)\right) - \left(b_1 e^{b_2 T} - \left(d_0 + \frac{b_1 b_2}{d_2} e^{(b_2 - d_2)T_{opt,A}^*} e^{d_2 T}\right)\right) \quad \text{(S13)}$$

Simplifying:

$$\mu_B(T) - \mu_A(T) = \left(\frac{b_1 b_2}{d_2} e^{(b_2 - d_2)T_{opt,A}^*} e^{d_2 T}\right) - \left(\frac{b_1 b_2}{d_2} e^{(b_2 - d_2)T_{opt,B}^*} e^{d_2 T}\right)$$

$$\mu_B(T) - \mu_A(T) = \frac{b_1 b_2}{d_2} e^{d_2 T}\left(e^{(b_2 - d_2)T_{opt,A}^*} - e^{(b_2 - d_2)T_{opt,B}^*}\right) \quad \text{(S14)}$$

Together, the assumptions that $b_2 < d_2$ and $T_{opt,A}^* < T_{opt,B}^*$ imply that

$$\left(e^{(b_2 - d_2)T_{opt,A}^*} - e^{(b_2 - d_2)T_{opt,B}^*}\right) > 0$$

Furthermore, this conclusion holds for any value of *T*. This means that, given the model expressed in equation (2) in the main text, a species with a higher $T_{opt}^*$ will also have a higher growth rate than all species with lower $T_{opt}$, across all temperatures. This is consistent with the approximation for $T_{min}$ (Equation S7), which suggests it is independent of $T_{opt}^*$, and $T_{max}$ (Equation S8), which increases linearly with $T_{opt}$. Together, these observations imply that niche width increases with $T_{opt}^*$. This scenario is both evolutionarily improbable (as it would lead to run-away selection for ever higher values of $T_{opt}^*$, a so-called "Darwinian demon") and inconsistent with prior empirical observations (Thomas et al. 2012, 2016). Consequently, it is likely that one or more of the other model parameters must be connected to $T_{opt}^*$, creating a constraint or tradeoff.

*S4.2 Potential constraints*

Many constraints are possible, affecting the shape and with of thermal reaction norms as optimum temperature increases. However, our approximations for the value of $T_{min}$ and $T_{max}$, motivate a few reasonable possibilities, all of which would cause $T_{min}$ to increase with $T_{opt}$. Specifically, $T_{min}$ depends on $d_0$ and $b_1$ while $T_{max}$ is independent of these values (recall, $T_{min} \approx \frac{ln(d_0/b_1)}{b_2} = \frac{ln(d_0)}{b_2} - \frac{ln(b_1)}{b_2}$ and $T_{max} \approx T_{opt}^* + \frac{\ln(d_2/b_2)}{d_2 - b_2}$). If $d_0$ increases or $b_1$ decreases (or both!) with increasing $T_{opt}^*$, the result would be an increase in $T_{min}$ with $T_{opt}^*$. This could occur in several ways:

*Hypotheses:*

(1) $b_1$ decreases with $T^*_{opt}$ (2) $d_0$ increases with $T^*_{opt}$ (3) $d_0/b_1$ increases with $T^*_{opt}$

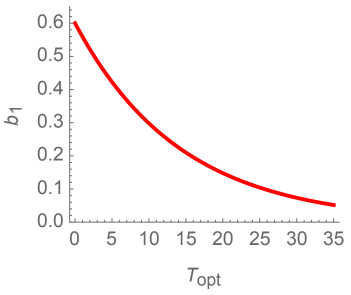


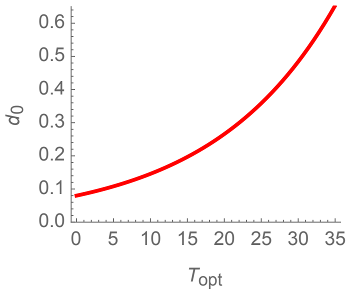


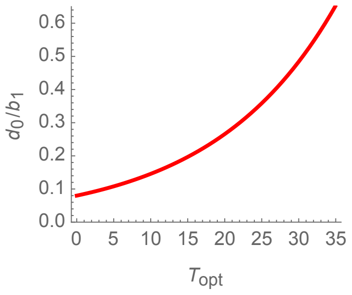


$$b_1 = a_0 e^{-a_1 T^*_{opt}} \qquad d_0 = a_0 e^{a_1 T^*_{opt}} \qquad d_0/b_1 = a_0 e^{a_1 T^*_{opt}}$$

$$\ln[b_1] = \ln[a_0] - a_1 T^*_{opt} \qquad \ln[d_0] = \ln[a_0] + a_1 T^*_{opt} \qquad \ln[d_0/b_1] = \ln[a_0] + a_1 T^*_{opt}$$

$$T_{min} \approx \frac{a_1}{b_2} T^*_{opt} + \frac{\ln[d_0/a_0]}{b_2} \qquad T_{min} \approx \frac{a_1}{b_2} T^*_{opt} + \frac{\ln[a_0/b_1]}{b_2} \qquad T_{min} \approx \frac{a_1}{b_2} T^*_{opt} + \frac{\ln[a_0]}{b_2}$$

In all cases, $T_{min}$ becomes approximately an increasing linear function of $T^*_{opt}$, with slope = $a_1/b_2$. If $a_1 > b_2$, then $T_{min}$ will increase more rapidly than $T_{max}$ as $T^*_{opt}$ increases, and niches will become narrower for species with higher optimum temperatures. Otherwise, if $a_1 < b_2$, species with higher optimum temperatures will have larger niche widths than species with lower optimum temperatures, although these thermal niches will still shift towards higher temperatures with $T^*_{opt}$. For example, assuming hypothesis (3) above, and manipulating $a_1$ vs $b_2$:

**$a_1 > b_2$**

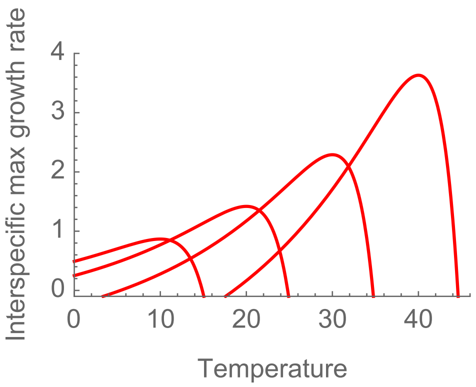


**$a_1 < b_2$**

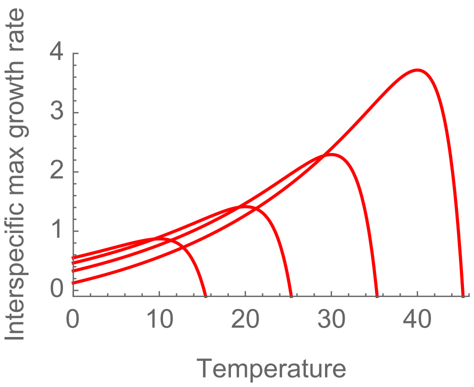

Referring to patterns of variation in the thermal traits of real species suggests that hypothesis (3) is most clearly supported by empirical evidence. It would result in a growth rate equation of

$$\mu(T,R) = b_1\left( e^{b_2 T} g(R) - \left(a_0 e^{a_1 T^*_{opt}} + \frac{b_2}{d_2} e^{(b_2-d_2)T^*_{opt}} e^{d_2 T}\right)\right) \tag{S15}$$

It is worth noting that this equation has an extra parameter (an expression with $a_0$ and $a_1$ replaced $d_0/b_1$, but $b_1$ still appears elsewhere in the equation). Fitting this equation to data on a single TPC is likely to cause problems, as it will not be possible to uniquely identify $a_0$ and $a_1$. Instead, we rely on estimating these parameters globally, by examining the relationship $\ln[d_0/b_1] = \ln[a_0] + a_1 T^*_{opt}$ across many curves, based on variation in estimated $d_0$, $b_1$, and $T^*_{opt}$ values (see main text and Appendix S3).

Section S5: Community envelopes

In certain contexts, it has proved useful to be able to approximate the upper limits of growth or function achieved across an entire suite of species that might inhabit a community, and how these limits change as a function of environmental conditions. A classic example of this is the Eppley curve (Eppley 1972), which described changes in maximum interspecific growth rate as a function of temperature (see also (Bissinger et al. 2008, Kremer et al. 2017). In certain circumstances, this function provides a reasonable approximation for the functioning of diverse communities (Norberg et al. 2001, Moisan et al. 2002, Thomas et al. 2012). We can derive similar envelope functions using the double exponential model (2) or its extensions (3) in the main text. This enables us to consider both which parameters drive the community-wide dependence of growth rate on temperature (of interest to MTE, etc.), and how this envelope changes when conditions deviate from optimal.

*S5.1 Deriving envelope functions*

First, consider $\mu(E_1, E_2, z)$, growth rate as a function of one (or more) environmental variables, $E_1$, $E_2$, …, and a trait $z$. The envelope providing an upper bound on $\mu$ across all possible trait values can be determined by finding the trait that maximizes $\mu$ at any given combination of environmental conditions. Specifically, we consider a function for $\mu$ that combines temperature and resource effects, while allowing the functional response $g(R)$ to be an arbitrary function that is independent of $T_{opt}^*$). We also assume that the $d_0/b_1$ vs. $T_{opt}^*$ tradeoff holds (see Equation S15), and then consider how growth rate changes as a function of species' trait $T_{opt}^*$

$$\frac{\partial\mu(T,N)}{\partial T_{opt}^*} = \frac{\partial}{\partial T_{opt}^*}\left(b_1\, e^{b_2 T} g(R) - \left(b_1 a_0 e^{a_1 T_{opt}^*} + \frac{b_1 b_2}{d_2} e^{(b_2-d_2)T_{opt}^*} e^{d_2 T}\right)\right)$$

$$= -b_1\left[a_0 \frac{\partial}{\partial T_{opt}^*}\left(e^{a_1 T_{opt}^*}\right) + \frac{b_2}{d_2} e^{d_2 T} \frac{\partial}{\partial T_{opt}^*}\left(e^{(b_2-d_2)T_{opt}^*}\right)\right]$$

$$= -b_1\left[a_0 a_1 e^{a_1 T_{opt}^*} + \frac{b_2(b_2-d_2)}{d_2} e^{d_2 T} e^{(b_2-d_2)T_{opt}^*}\right] \qquad \text{(S16)}$$

Note that this derivative is independent of any resource effects; consequently, the optimal $T_{opt}^*$ value will be independent from resource levels. The next step is to solve this equation for the specific $T_{opt}^*$ value $\hat{T}_{opt}$ that satisfies the equation $\left.\frac{\partial\mu(T,N)}{\partial T_{opt}^*}\right|_{T_{opt}^*=\hat{T}_{opt}} = 0$. The result is

$$\hat{T}_{opt} = \frac{1}{a_1 + d_2 - b_2}\left(d_2 T + \ln\left[\frac{b_2}{a_0 a_1}\left(1 - \frac{b_2}{d_2}\right)\right]\right) \qquad \text{(S17)}$$

The final step is to substitute Equation S17 into Equation S15. The result is a general equation for the maximum growth rate achievable (by manipulating $T_{opt}^*$) as a function of temperature ($T$) and resource ($R$),

$$\mu_{comm}(T,R) = b_1\left(e^{b_2 T} g(R) - \left(a_0 e^{a_1 \hat{T}_{opt}} + \frac{b_2}{d_2} e^{(b_2-d_2)\hat{T}_{opt}} e^{d_2 T}\right)\right) \qquad \text{(S18)}$$

Note that the resource limitation term, $g(R)$, has returned, indicating that resource limitation will affect the height of this envelope function, but not the optimal $\hat{T}_{opt}$ trait at a given temperature.

*S5.2 Simplifying envelope function*

We can obtain additional insights from Equation S18. In particular, we seek to clarify the effect of temperature on the scaling of maximum growth rate across taxa. To begin, let

$$C_1 = \ln\left[\frac{b_2}{a_0 a_1}\left(1 - \frac{b_2}{d_2}\right)\right]$$

$$C_2 = \frac{1}{a_1 + d_2 - b_2}$$

This simplifies the $\hat{T}_{opt}$ term in Equation S18 for $\mu_{comm}$ to

$$\mu_{comm}(T,R) = b_1\left(e^{b_2 T} g(R) - \left(a_0 e^{a_1 C_2 (d_2 T + C_1)} + \frac{b_2}{d_2} e^{d_2 T + C_2(b_2 - d_2)(d_2 T + C_1)}\right)\right)$$

Expanding exponential terms and re-arranging yields

$$\mu_{comm}(T,R) = b_1\left(e^{b_2 T} g(R) - a_0 e^{a_1 C_1 C_2} e^{a_1 C_2 d_2 T} + \frac{b_2}{d_2} e^{C_1 C_2 (b_2 - d_2)} e^{d_2 T\left(1 + C_2(b_2 - d_2)\right)}\right) \quad \text{(S19)}$$

Next, we observe that algebraic manipulation of the definition of $C_2$ yields

$$a_1 C_2 = 1 + C_2(b_2 - d_2)$$

Substituting this into Equation S19 and simplifying, we obtain

$$\mu_{comm}(T,R) = b_1 e^{b_2 T} g(R) - b_1\left(a_0 e^{a_1 C_1 C_2} + \frac{b_2}{d_2} e^{C_1 C_2 (b_2 - d_2)}\right) e^{a_1 C_2 d_2 T}$$

Next, we define two more constants, noting that they are independent of temperature *T*:

$$C_3 = a_1 C_2 \quad \text{(S20)}$$

$$C_4 = b_1\left(a_0 e^{a_1 C_1 C_2} + \frac{b_2}{d_2} e^{C_1 C_2 (b_2 - d_2)}\right) \quad \text{(S21)}$$

Substituting and simplifying produces

$$\mu_{comm}(T,R) = b_1 e^{b_2 T} g(R) - C_4 e^{C_3 d_2 T} \tag{S22}$$

This is the derivation of equation (6) in the main text.

Recall that the purpose of this analysis was to find an expression for the envelope constraining maximum growth rate (across all species in a community) as a function of temperature. Equation S22 provides this result in a simplified form that highlights the fact that the community-level envelope itself is determined by the difference between two exponential functions of temperature. In this way, it is fundamentally similar to the definition of the population-level double exponential model (1), including the potential effects of resource limitation, albeit with different parameters associated with the loss term.

*S5.3 Approximating the behavior of the community envelope function*

One of our additional goals is to understand whether the community envelope emerging from this analysis is consistent with empirical work suggesting that (i) maximum growth rate across species increases exponentially with temperature (the Eppley curve), and (ii) the scaling of this increase is consistent with the temperature dependence of birth or photosynthesis (governed by $b_2$ in our model). To help consider this, we observe that Equation S22 can be further re-arranged. If we define $C_5 = \frac{(a_1 - b_2)}{(1 + a_1/(d_2 - b_2))}$, then Equation S22 can be manipulated as follows:

$$\mu_{comm}(T,R) = b_1\, e^{b_2 T}(g(R) - C_4\, e^{C_3\, d_2\, T - b_2\, T})$$

$$= b_1\, e^{b_2 T}\left(g(R) - (C_4/b_1)\, e^{\frac{(a_1 - b_2)}{(1 + a_1/(d_2 - b_2))}T}\right)$$

$$= b_1\, e^{b_2 T}\left(g(R) - \left(\frac{C_4}{b_1}\right)\, e^{C_5 T}\right) \tag{S23}$$

Now, we note that Equation S23 will be similar to a basic exponential function of temperature $T$ if $(C_4/b_1)$ is very close to zero, if $C_5$ is $\leq 0$, or both.

*Case 1:* $(C_4/b_1)$ *is very close to zero*. As defined in Equation S21, $C_4$ must be positive. However, if it is a sufficiently small number close to zero, then the effects of the second exponential term in Equation S23, $e^{C_5 T}$, will be minimal across biologically realistic temperature ranges.

Subsequent cases depend on the quantity $C_5$, which in turn depends on two critical terms, $(a_1 - b_2)$ and $(d_2 - b_2)$.

*Case 2:* $a_1 = b_2$. This results in $C_5 = 0$. Then, our envelope function collapses exactly into an exponential function of $b_2$ and temperature, recapitulating the Eppley curve, albeit with a modified intercept.

*Case 3:* $a_1 < b_2$. Then $C_5 < 0$, and although the term $C_4\ e^{C_5 T}$ will depend on temperature, it will become increasingly small and irrelevant as $T$ increases, and as $a_1 \ll b_2$. Consequently, we expect that overall Equation S23 would behave nearly exponentially over reasonable temperature ranges.

*Case 4:* $a_1 > b_2$, *but only slightly*. Then $C_5$ will be near zero, and again Equation S23 will be nearly exponential.

*Case 5:* $a_1 \gg b_2$. Then $e^{C_5 T}$ will be large and Equation S23 will deviate from a simple exponential.

*Case 6:* $(d_2 - b_2) < a_1$. By definition, $a_1$ and also $(d_2 - b_2)$ will always be positive (as we require $d_2 > b_2$) so the denominator of $C_5$ will always be positive. If $d_2 - b_2$ is very small (relative to $a_1$), then again $C_5$ will approach 0, and Equation S23 will converge on an exponential expression.

Invoking realistic, empirical parameters for autotrophs, additional numerical analyses reveal that Equation S23 is nearly indistinguishable from a simple exponential function over biologically relevant ranges of temperature (here, 0-30°C) (see Appendix S2: Figure S1 below). These conclusions rest on considering the integral of the squared difference between Equation S23 and a generic exponential function of temperature, $f(T) = \phi_1 e^{\phi_2 T}$, or

$$\int_0^{30} \left(f(T) - (b_1(g(R) - (C_4/b_1)\, e^{C_5 T})\, e^{b_2 T})\right)^2 dT$$

then solving for the values of $\phi_1$ and $\phi_2$ that minimize this quantity. In effect, this equivalent to using a nonlinear least squares approach to fitting a regression, but fitting one continuous function to another function rather than to observational data. While it is not possible to analytically solve for $\phi_1$ and $\phi_2$, numerical methods can provide a solution given particular selections of parameters.

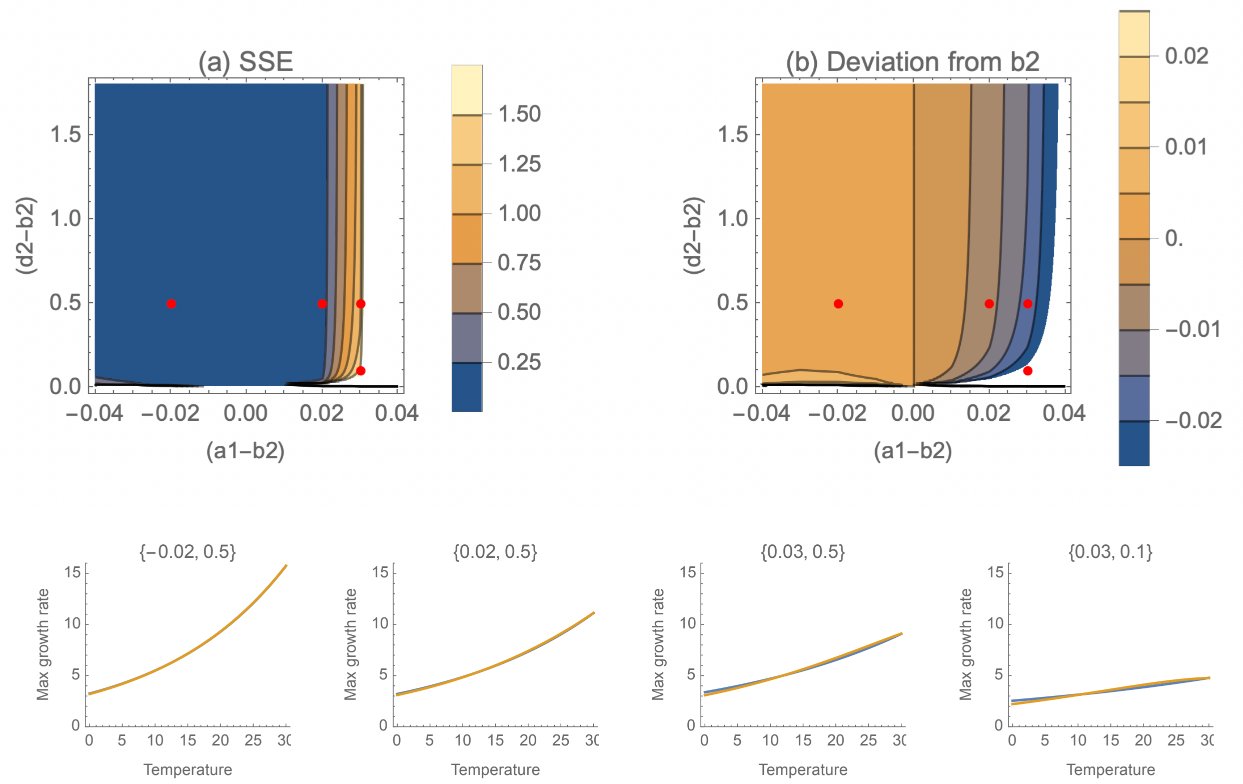


**Figure S1**. We can use a simple exponential function to approximate the actual envelope function (Equation S19) over the range 0-30°C, assuming $\ln(a_0) = -1.82$ (see Appendix S1: Table S5 for autotrophs), $b_1 = 3.97$ (median value across all autotrophic curves), and $b_2 = 0.0498$ (MTE value for autotrophs). This numerical approximation is accomplished by minimizing the summed squared errors between Equation S23 and an exponential function. The summed squared error of the resulting fit (a) and the difference between the estimated exponent/thermal sensitivity and $b_2$ (b) can then be calculated assuming a range of values for $d_2$ and $a_1$ (expressed relative to $b_2$). Note that the approximation is exact when $a_1 = b_2$. Empirically, using the best available data, the 95% confidence interval surrounding the best estimate of $b_2$ (Eppley coefficient) spans roughly ±0.01 (Kremer et al. 2017); this provides a benchmark for assessing the values in (b). Generally, the approximation is extremely good when $a_1 < b_2$ across a wide range of $d_2$ values. The approximation also performs well when $(a_1 - b_2) < 0.03$, except for values of $d_2$ close to $b_2$. Finally, the bottom row of figures shows examples of Equation S23 (blue) and the simple exponential approximation (orange) for various locations in (a) and (b) (red dots, panels go from left to right, then down). These illustrate that the exponential approximation is extremely close; however, it's important to distinguish between whether or not the exponential approximation fits Equation S23 well (all four examples), and whether the resulting exponential temperature sensitivity recapitulates the temperature scaling implied by $b_2$ (only the first 1-3 examples), including specific predictions of the MTE.

Our empirical investigations suggest that both autotrophs and heterotrophs likely fall into Cases 2 or possibly case 4, $b_2 \leq a_1$, (Appendix S1: Table S7), and consequently should have limiting envelopes that are nearly or exactly exponential functions of temperature with a scaling exponent quite close to $b_2$. Together, these observations suggest that – when resources are not limiting – the overall maximum interspecific growth rate should scale with temperature as an exponential function with a rate that is quite similar to $b_2$. It follows that if $b_2$ – the temperature dependence of birth rate – can be interpreted as the sensitivity of photosynthesis to temperature (see main text), the predicted envelope function is compatible with developments harmonizing the Eppley curve with Metabolic Theory (Kremer et al. 2017).

*S5.4 Comments on the effects of resource limitation*

In addition to this connection to established theoretical predictions, Equation S23 also offers insights into how resource limitation might affect the sensitivity of interspecific maximum growth rates to temperature. As resource limitation kicks in, the term $g(R)$ generally converges on zero for a range of typical models and resources (including Monod resource dynamics, or irradiance curves such as Eilers-Peeters). In the process, there will be a region where $g(R) \approx (C_4/b_1)\, e^{C_5 T}$. When this occurs the relationship between maximum interspecific growth rate and temperature may appear dampened or even flat over a biologically relevant range of temperatures. This is compatible with the results of prior studies such as that of (Edwards et al. 2016), which found that light limitation weakens the temperature dependence of maximum growth rate across species (see main text).
