## Appendix 3: Data filtering and quality control for "How interactions between temperature and resources scale from populations to communities"

### Appendix S3: Filtering and quality control for microbial TPCs

Previously published compilations of experimental data on the effects of temperature on microbial growth rate (Thomas et al. 2016, Corkrey et al. 2016) provide the raw information for our study of variation in the parameters of the double exponential model. Prior to this investigation, however, these data sets required additional filtering and quality control to ensure that the analyses are appropriate and meaningful.

From the Corkery et al. (2016) data set we dropped taxa from the Animalia and Fungi kingdoms (while retaining members of Archaea, Eukaryota, and Bacteria), excluded non-heterotrophs (as there was significant overlap with data from Thomas et al. 2016), and removed strains 900 and 903, which appeared to be exact duplicates of other strains. For the Thomas et al. (2016) autotrophic data set some additional experimental conditions – aside from temperature – are reported, including light, day length, salinity, and nutrient concentrations. Where available, we used these conditions to exclude experimental results where conditions were sub-optimal due to factors other than temperature. In this step, we followed criteria used previously with this data set (Thomas et al. 2012, Thomas et al. 2016); full details are also provided in the associated R code. Finally, for both data sets, we also removed strains with growth rates measured at fewer than five distinct temperatures, and thermophiles (which we defined as strains studied using average experimental temperatures exceeding 35°C). After all of these exclusions, we were left with 271 and 363 individual strains for heterotrophs and photoautotrophs, respectively.

Because our interest is in fully capturing unimodal TPCs, we also decided to exclude cases where only ascending or descending portions of a TPC are documented. To do this in an automated fashion, we fit both the double exponential model (2) and a simpler model that only allows growth rates to increase or decrease exponentially with temperature to data for each strain. When the simpler model had a lower AIC value than the double exponential model (by >2AIC units), we excluded the corresponding strain's TPC from all further analyses; these reflect situations where the data are insufficient to rigorously characterize a unimodal growth curve. This reduced the totals to 251 and 321 heterotrophic and photoautotrophic curves, respectively.

We also removed all remaining curves where the double exponential model on its own had an $R^2$ <= 0.85 (a loss of ~11% of our remaining strains). To ensure that the curve fits were adequate to characterize both the upper and lower thermal limits for each TPC, we also used estimates of $T_{min}$ and $T_{max}$ as filtering criteria. Because these traits are not explicit parameters of main text equation (2) (see Appendix S2: Section 2) they were instead calculated using numerical methods (see code in Appendix 4). Consistent with prior publications, we excluded cases where $T_{min}$ and/or $T_{max}$ either could not be calculated or where their inferred values fell farther than 5°C from the closest experimental temperature treatments (to avoid error due to extrapolation). This reduced the data set by ~50%. The remaining data included a total of 248 thermal reaction norms characterized reliably by the double exponential model, 113 heterotrophs and 135 photoautotrophs.

Regarding trophic mode, we noted that the Thomas et al. (2016) data contain both cryptophytes and dinoflagellates; members of these groups are known to often exhibit mixotrophy, engaging in both photosynthesis and heterotrophic consumption of bacteria and other microbes, often depending on environmental conditions. To more directly test our hypotheses (regarding differences in temperature dependences between heterotrophs and photoautotrophs), we exclude these potentially mixotrophic taxa (accounting for 25 of the photoautotrophs) from further analyses, although the associate data are made available.
