## Appendix 4: Guide to code and supporting analyses for "How interactions between temperature and resources scale from populations to communities"

**Appendix S4: Guide to code supporting analysis and results**

**Table S1:** All code and data associated with the manuscript's analysis and visualizations are made available by the lead author on figshare. Data are here (XXXX) and code is here (XXXX). Both will be made publicly available following publication. For convenience, this table provides a summary each code files including their purpose, inputs, and outputs, so readers can easily navigate to the relevant materials.

| File | Purpose | Inputs | Outputs |
|---|---|---|---|
| double_exponential_model_math.nb | Mathematical analysis of the double exponential model | | Fig. 1<br>Fig. 2 a,b,d<br>Fig. 4 a-c,<br>Appendix S1: Fig. S2-S4<br>Appendix S2: Fig. S1 |
| resource_limitation_and_temperature_traits.R | Data processing and analysis of effects of nutrients, light on optimum temperature | Bestion_Growth rate data.csv<br>gcb13641-sup-0002-appendixs1.csv<br>lno10282-sup-0005-supptables.xlsx | Fig. 2c,e<br>Appendix S1: Table S2, S3<br>Topt_x_N_derived_data.csv |
| TPC_database_processing.R | To clean up, organize and combine data on microbial growth rates at different temperatures | curve data_20140606.csv<br>isolate data_20140606.csv<br>growth data_20140606.csv | TxNxL_thinned_curve_derived_data.csv<br>(a quality-controlled subset of raw growth curves for phytoplankton) |
| double_exponential_fitting.R | To fit the double exponential model (2) to the quality controlled | TxNxL_thinned_curve_derived_data.csv<br>isolate data_20140606.csv | double_exponential_trait_ests_QCd_derived_data.csv |

| | | | |
|---|---|---|---|
| | microbial growth rate data | | |
| double_exponential_model_stats_analyses.R | Analysis of the trait estimates produced by fitting the double exponential model to data | double_exponential_trait_ests_QCd_derived_data.csv | Final analyses and figures, Fig. 3<br>Appendix S1: Tables S4-S7, Figure S1 |
